# A nuclear receptor facilitates differentiation of human PSCs into more mature hepatocytes

**DOI:** 10.1101/2022.03.24.485641

**Authors:** Haiting Ma, Esmée de Zwaan, Yang Eric Guo, Paloma Cejas, Prathapan Thiru, Martijn van de Bunt, Jacob F. Jeppesen, Sudeepa Syamala, Alessandra Dall’Agnese, Brian J. Abraham, Dongdong Fu, Carrie Garrett-Engele, Tony Lee, Henry W Long, Linda G. Griffith, Richard A. Young, Rudolf Jaenisch

## Abstract

The capacity to generate functional hepatocytes from renewable human pluripotent stem cells (hPSCs) could address limited supplies of primary human hepatocytes. However, hepatocytes differentiated from hPSCs *in vitro* are functionally immature. To understand mechanisms regulating maturation of *in vitro* derived hepatocytes, we developed a 3D spheroid differentiation system and compared gene regulatory elements in uncultured human primary hepatocytes with those in hepatocytes that were differentiated in 2D or 3D conditions from human PSCs by RNA-seq, ATAC-seq, and H3K27Ac ChIP-seq. Three-dimensional differentiation improved enhancer activity and expression of transcription factor *ONECUT1*, but was insufficient to upregulate human-specific mature hepatocytes marker gene *CYP3A4* or super-enhancer regulated transcription factor gene *NFIC*. Regulome comparisons showed reduced enrichment of thyroid receptor THRB motifs in accessible chromatin and in active enhancers without reduced transcription of *THRB*, suggesting the regulation at the level of THRB ligands in PSC-differentiated hepatocytes. Addition of thyroid hormone T3 to the PSC-differentiated hepatocytes increased *CYP3A4* expression. T3 increased binding of THRB to the *CYP3A4* proximal enhancer and restored the super-enhancer status and gene expression of *NFIC* and reduced expression of *AFP*. The resultant hPSC-hepatocytes showed gene expression, epigenetic status and super-enhancer landscape closer to primary hepatocytes and activated regulatory regions including non-coding SNPs associated with liver-related diseases. Transplanting the 3D PSC-hepatocytes into immunocompromised mice resulted in engraftment of human hepatocytes in the mouse liver parenchyma without disrupting normal liver histology at 6 months after transplantation. This work provides insights into the functions of nuclear receptor THRB and highlights the importance of the environmental factors-nuclear receptors axis in regulating maturation of human PSC-differentiated cell types.

## Introduction

The liver is an essential organ with diverse functions including digestion, metabolism, and detoxification (Godoy et al., 2013) with hepatocytes being the major parenchymal cell type carrying out the essential functions (Aizarani et al., 2019; MacParland et al., 2018). For example, hepatocytes absorb multiple chemicals including many clinically used drugs by drug transporter proteins (Fahrmayr et al., 2010) and metabolize the chemicals by cytochrome P450 enzymes including CYP3A4 and CYP2C9 (Zanger and Schwab, 2013). Furthermore, liver or hepatocyte transplantations are the only known effective treatments for multiple end-stage liver disease (Bhatia et al., 2014). Due to limitations of alternative sources, primary hepatocytes remain the gold standard for hepatocytes, presenting a challenge because of limited supply and of variability associated with genetic and environmental characteristics of donors. Facing the demand for hepatocytes for research and clinical uses, generating functional and mature hepatocytes from sources including *in vitro* expansion of primary hepatocytes (Hu et al., 2018; Michailidis et al., 2020; Peng et al., 2018), adult stem cells (Huch et al., 2015), human fibroblast (Xie et al., 2019) and renewable human pluripotent stem cells have been recognized as possible solutions (Zaret and Grompe, 2008).

Mechanisms regulating hepatocytes specification during embryonic development (Prior et al., 2019; Si-Tayeb et al., 2010b) provided insights for *in vitro* differentiation of hepatocyte-like cells from hPSC with 2D differentiation protocols (Ang et al., 2018; Si-Tayeb et al., 2010a). Although various 3D differentiation systems (Chen et al., 2020; Ogawa et al., 2013; Takebe et al., 2013) improved hepatocyte gene expression, the *in vitro* generated hepatocyte-like cells showed function and gene expression profiles similar to fetal hepatocytes (Camp et al., 2017; Chen et al., 2018). The mechanisms resulting in improved gene expression with 3D differentiation and the epigenetic difference between PSC-hepatocytes and primary hepatocytes *in vivo* remain to be characterized. *In vivo* environmental factors including hormones, nutrients, and metabolites that are different from the *in vitro* culture conditions could contribute to functional maturation by epigenetic mechanisms (Chen et al., 2018).

We developed a 3D spheroid hepatocyte differentiation system, and profiled gene expression, chromatin accessibility, and enhancer landscapes of uncultured primary human hepatocytes and PSC-differentiated hepatocytes in 2D and 3D differentiation. In contrast to 2D differentiated hepatocytes, 3D differentiation improved expression of *ONECUT1*, a critical liver enriched transcription factor (Kyrmizi et al., 2006) but expression of CYP genes including *CYP3A4* remained low. Compared to primary hepatocytes, 3D differentiated hepatocytes showed reduced activity for nuclear receptor THRB. Addition of the THRB ligand T3 increased *CYP3A4* expression and improved both gene expression and H3K27Ac status for multiple hepatocyte genes, including the liver super-enhancer-regulated gene *NFIC*. These results suggest that environmental influences relayed by nuclear receptors regulate cell states in the process of hepatocyte maturation and the cell maturation states can be modulated by their ligands.

## Results

### THRB motif enrichment in accessible chromatin and active enhancers differed between primary human hepatocytes and PSC-hepatocytes

We differentiated H1-OCT4GFP hPSCs (Zwaka and Thomson, 2003) into definitive endoderm(Ma et al., 2020), followed by liver induction with FGF2 and BMP4 (Si-Tayeb et al., 2010b), and William’s E based medium to generate hepatocyte-like cells with improved ALB expression, reduced AFP expression, and similar proportions of cells expressing HNF4A (**Figure S1**). Based on the 2D differentiation protocol, we developed a spheroid-based 3D hepatocyte differentiation system (**Figure 1A**) with the following modifications. 1. Adding a gut tube stage by FGF7 treatment after definitive endoderm stage (**Figure S1E**). 2. Including a Notch inhibitor and a TGFβ inhibitor to reduce cystic formation that suggests bile duct epithelial cells(Huch et al., 2015) (**Figure S1F**). 3. Dissociated and aggregated cells to spheroids during stage 3 (**Figure S1G**). 4. Using less insulin (1%) in the culture medium, and adding forskolin which was shown to maintain primary hepatocytes in long-term culture (Xiang et al., 2019) and to improve hepatocyte differentiation (Ogawa et al., 2013). Additionally, we supplemented the medium with linoleic acid and micronutrients. The 3D differentiation protocol (**Figure 1A**) generated hepatocyte-like cells with improved expression of hepatocytes markers compared to 2D differentiation, although fetal hepatocyte gene *AFP* remained highly expressed (**Figure S1I-J)**.

**Fig. 1.**
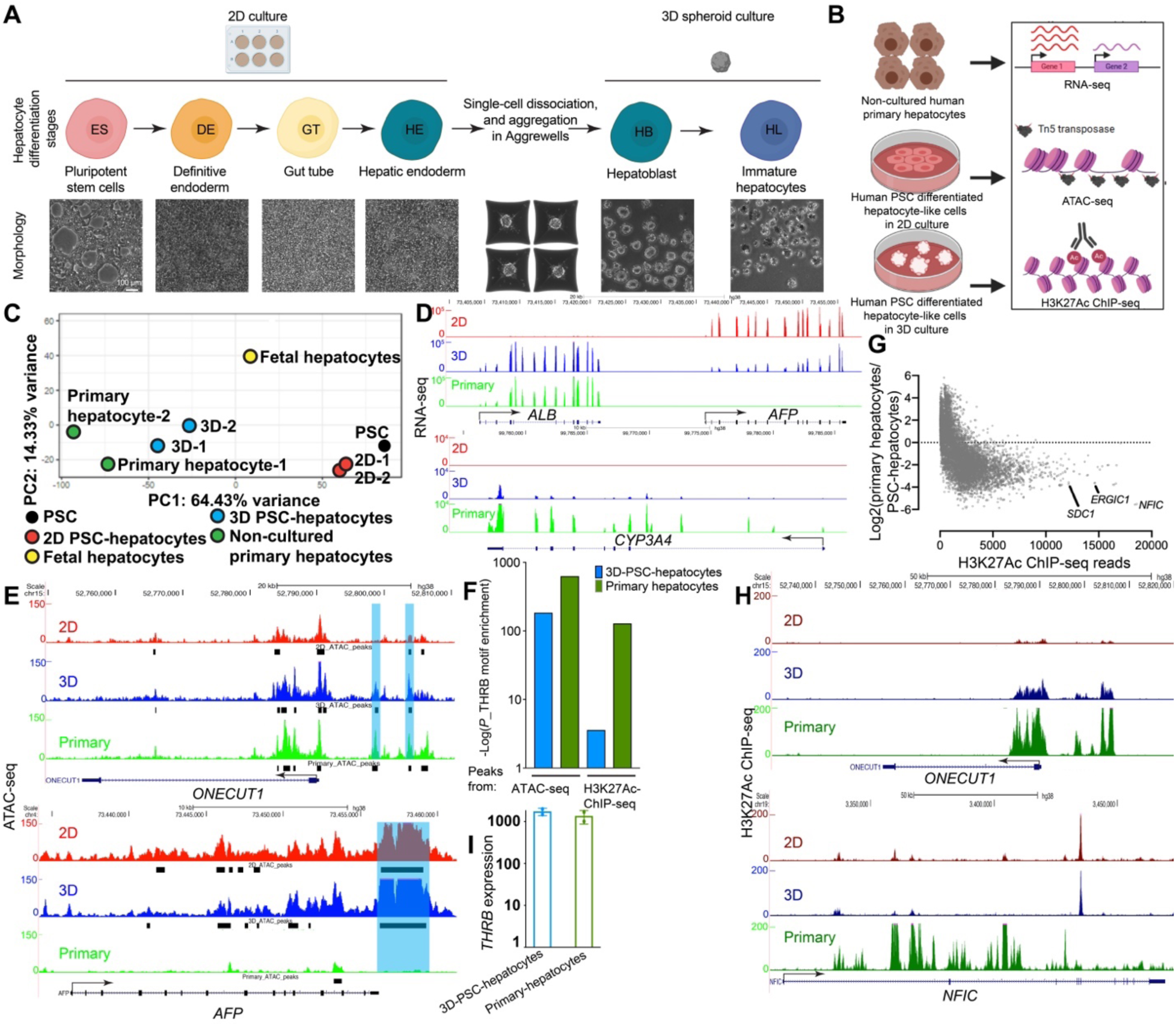
**Regulome analysis identified differences in THRB motif enrichment in accessible DNA and active enhancers between primary hepatocytes and PSC-differentiated hepatocytes.** (A) Development of a spheroid-based hepatocytes differentiation system from hPSC. (B) A cartoon diagram of experimental design. Uncultured primary hepatocytes, 2D PSC-hepatocytes, and 3D PSC-hepatocytes were subjected to RNA-seq, ATAC-seq, and H3K27Ac ChIP-seq. (C) Principal component analysis (PCA) of 2D hPSC-hepatocytes, 3D hPSC-hepatocytes, uncultured primary hepatocytes, and fetal hepatocytes. (D) Genome browser gene tracks representing RNA-seq results of the *ALB*/*AFP* and *CYP3A4* loci from 2D PSC-hepatocytes (2D), 3D PSC-hepatocytes (3D), and primary hepatocytes (Primary). (E) ATAC-seq tracks of *ONECUT1* and *AFP* loci from 2D PSC-hepatocytes (2D), 3D PSC-hepatocytes (3D), and primary hepatocytes (3D). (F) Enrichment of THRB motifs (-log*P*) in the ATAC-seq peaks and H3K27Ac ChIP-seq peaks in 3D-PSC-hepatocytes (blue) and primary hepatocytes (green). (G) Differential analysis of H3K27Ac ChIP-seq between primary hepatocytes and PSC-hepatocytes. (H) H3K27Ac ChIP-seq tracks of *ONECUT1* and *NFIC* loci from 2D PSC-hepatocytes (2D), 3D PSC-hepatocytes (3D), and primary hepatocytes (3D). (I) Expression of *THRB* in 3D PSC-hepatocytes and primary hepatocytes. Data showing mean ± standard error, *n*=2. The scale bar in (A): 100 µm.

To understand the transcriptional basis of immaturity of PSC-hepatocytes, we compared mRNA-transcriptomes of non-cultured human primary hepatocytes and hPSC differentiated hepatocytes under 2D and 3D conditions (**Figure 1B**). Principal component analyses of RNA-seq results showed that 3D PSC-hepatocytes grouped closer with primary hepatocytes, whereas 2D PSC-hepatocytes were closer to fetal hepatocytes (RNA-seq results from (Xie et al., 2019)) (**Figure 1C**). KEGG pathway analysis identified several hepatocyte functions (including complement, coagulation factors, and cytochrome P450) were downregulated in 2D PSC-hepatocytes compared to primary hepatocyte (**Figure S2B, *P*<0.05**). Gene set enrichment analysis (GSEA) also showed hepatocyte functions such as the drug metabolism/CYP P450 pathways and protein secretion were downregulated, whereas cell cycle regulators and receptors-ECM interactions were upregulated in 2D PSC-hepatocytes as compared to primary hepatocytes (**Figure S2 C-D, nominal *P*-value<0.05**). In contrast, these components were not significantly different between 3D PSC-hepatocytes and primary hepatocytes (**Figure S2F-G**). Compared to 2D PSC-hepatocytes, 3D PSC-hepatocytes showed improved expression of genes associated with liver functions (**Figure S2 H-J**). Consistent with gene expression analysis (**Figure S1J**), *ALB* expression was increased about 9-fold in 3D as compared to 2D PSC-hepatocytes (**Figure 1D, Figure S1J**). The expression of a key drug metabolism enzyme and mature hepatocyte marker *CYP3A4* also increased about 80-fold in 3D as compared to 2D PSC-hepatocytes (**Figure 1D, Figure S1J**). However, the absolute expression levels were only approximately 1% of those in primary hepatocytes. We conclude that culturing PSC-hepatocytes in 3D increases expression of complements and coagulation factors but failed to improve hepatocyte maturation genes including *CYP3A4*.

To examine gene regulatory elements that are typically located in accessible chromatin regions, we performed ATAC-seq (Buenrostro et al., 2013) on 2D PSC-hepatocytes, 3D PSC-hepatocytes, and primary hepatocytes (**Figure 1B**). Consistent with improved expression of *ONECUT1* in 3D PSC-hepatocytes compared to the 2D PSC-hepatocytes (**Figure S1J**), 3D PSC-hepatocytes showed increased chromatin accessibility at the 5’ region of *ONECUT1* locus (**Figure 1E, top panel**). In contrast, the *AFP* gene that was expressed in PSC-hepatocytes showed ectopic chromatin accessibility at the 3’ of *AFP* locus as compared to primary adult hepatocytes (**Figure 1E, bottom panel**). Transcription factor motifs enrichment analysis in ATAC-seq peaks showed more significant enrichment of the thyroid hormone receptor β (THRB) motif in primary hepatocytes as compared to 3D PSC-hepatocytes (**Figure 1F**).

We also performed H3K27Ac ChIP-seq to survey active promoters and enhancers landscape(Creyghton et al., 2010) in the three cell types (**Figure 1B**). Comparative analyses of primary hepatocytes and PSC-hepatocytes indicated lower enhancer activity in PSC-hepatocytes as compared to primary hepatocytes (**Figure 1G**). Consistent with the gene expression and chromatin accessibility data, 3D differentiation improved H3K27Ac signal of the *ONECUT1* locus (**Figure 1H**). Furthermore, we identified super-enhancers associated genes, a group of genes with of disproportionally high density of H3K27Ac ChIP-seq signals (Whyte et al., 2013). *NFIC* was identified as one of the top super-enhancer genes in primary hepatocytes and showed increased H3K27Ac signals compared to PSC-hepatocytes (**Figure 1H**). Additionally, correlation studies of super-enhancer and RNA-seq results showed super-enhancer tended to be higher expressed (**Figure 2A-E**). Analysis of super-enhancer genes from H3K27Ac ChIP-seq and highly expressed genes from RNA seq showed two groups of genes: super-enhancer associated genes that are more regulated and are associated with disease (insulin resistance, type 2 diabetes), whereas highly expressed but non-super-enhancer-regulated genes were enriched in more basic hepatocyte functions such as nitrogen metabolism (**Figure 2F**).

**Figure. 2.**
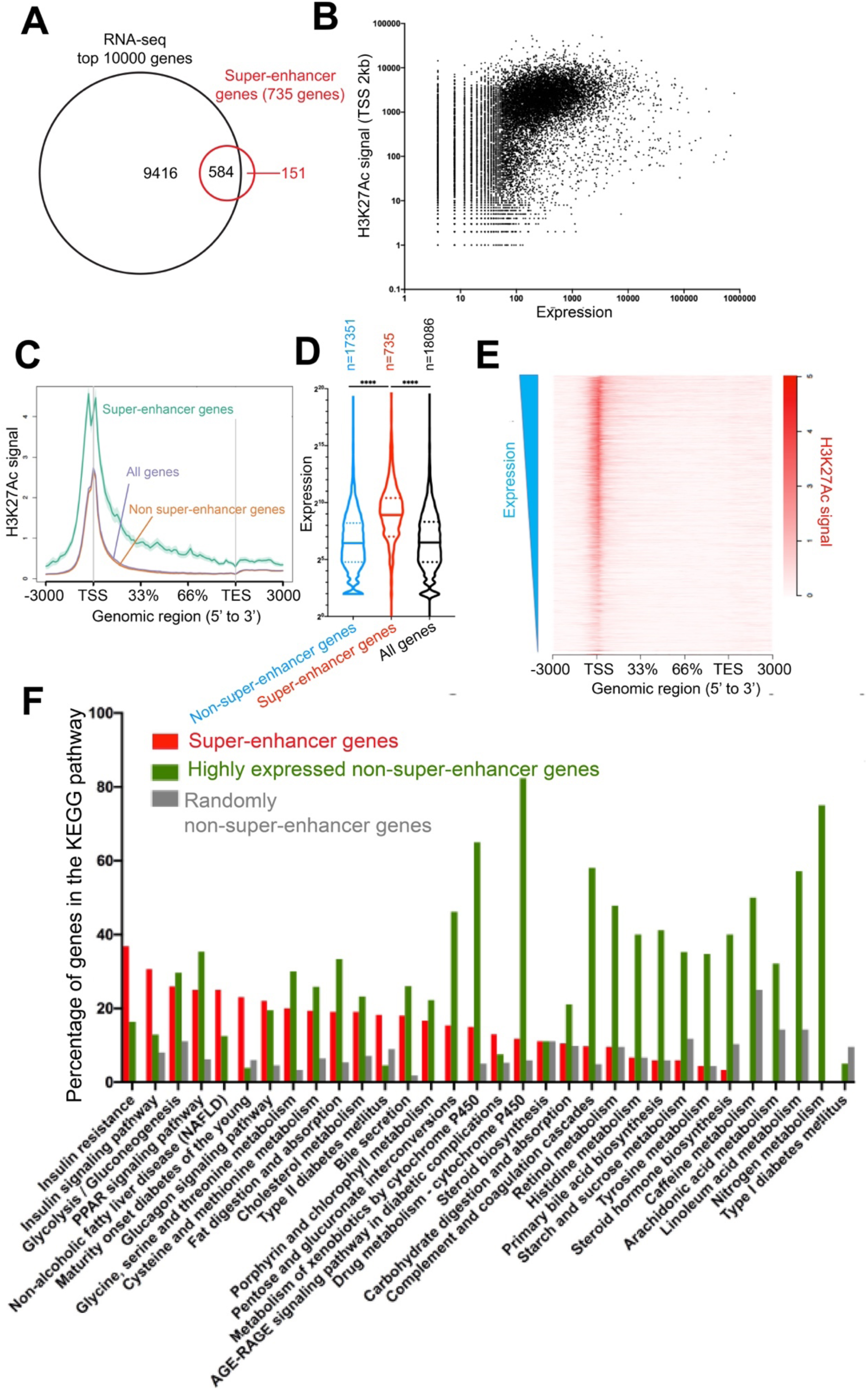
**Correlation between gene expression and H3K27Ac ChIP-seq signals.** (A) Overlapping between super-enhancer genes and top 10000 expressed genes from RNA-seq from primary hepatocytes. (B-E) Correlation between gene expression and H3K27Ac ChIP-seq signals. (F) Super-enhancer genes list was used to query KEGG database, and proportion of super-enhancer genes in the pathways were plotted (red bars). A list of highly expressed genes that are not super-enhancer genes were subjected to the same analysis and proportion of highly expressed non-super-enhancer genes in the pathways were plotted (green bars). Randomly selected genes were used as a control (grey bars).

Motif analysis from the H3K27Ac ChIP-seq showed that THRB motifs were more significantly enriched in primary hepatocytes than in 3D PSC-hepatocytes (**Figure. 1F**). However, gene expression of *THRB* did not correlate with the reduced enrichment of THRB motifs in accessible chromatin and active enhancers (**Figure 1I**). Transcription factor THRB is a nuclear receptor (NR1A2) that form heterodimeric DNA binding complex with RXRA based on availability of cognate ligands, thereby relaying information from the environment to the cells (Evans and Mangelsdorf, 2014). These data suggest that the differential enrichment of THRB motifs in accessible chromatin and active enhancers between primary hepatocytes and 3D-PSC-hepatocytes are potentially regulated at the levels of THRB ligand availability.

### THRB upregulated *CYP3A4* by binding to the *CYP3A4* proximal enhancer

We hypothesized that THRB may improve PSC-hepatocytes maturity. Addition of 3 µM T3 to the medium increased *CYP3A4* transcript (**Figure 3A**) and protein (**Figure 3B**) levels. In contrast, adding retinoic acid (RA, the ligand for another nuclear receptor RARA) did not improve *CYP3A4* expression (**Figure S3A**). Furthermore, T3 was required for the maintenance of the high expression of *CYP3A4* (**Figure S3B**). To examine the mechanisms of upregulation of *CYP3A4* by T3, we identified a promoter and two 5’ enhancers (proximal and distal enhancer) active in primary hepatocytes and 3D PSC-hepatocytes (**Figure 3C**). Analysis of *CYP3A4* 5’ regulatory elements determined that 3D PSC-hepatocytes with lower *CYP3A4*expression compared to primary hepatocytes, had weak signals at the proximal enhancer compared to primary hepatocytes. Furthermore, a THRB binding motif was detected in the proximal enhancer that overlapped with an ATAC-seq peak in 3D PSC-hepatocytes (**Figure 3C**). To test if THRB could bind to the proximal enhancer, we examined anti-FLAG ChIP-seq datasets from HepG2 cells with a C-terminal FLAG tagging of the endogenous *THRB* gene (THRB-FLAG HepG2) (Partridge et al., 2020) and found anti-FLAG-THRB ChIP-seq peaks overlapping with the proximal enhancer (**Figure 3C**). **Figure 3D** shows that in primary hepatocytes 74% and in 3D PSC-hepatocytes 54% of the ATAC-seq peaks overlapped with THRB peaks, consistent with contribution of THRB binding to gene regulatory elements in mature hepatocytes.

**Figure. 3.**
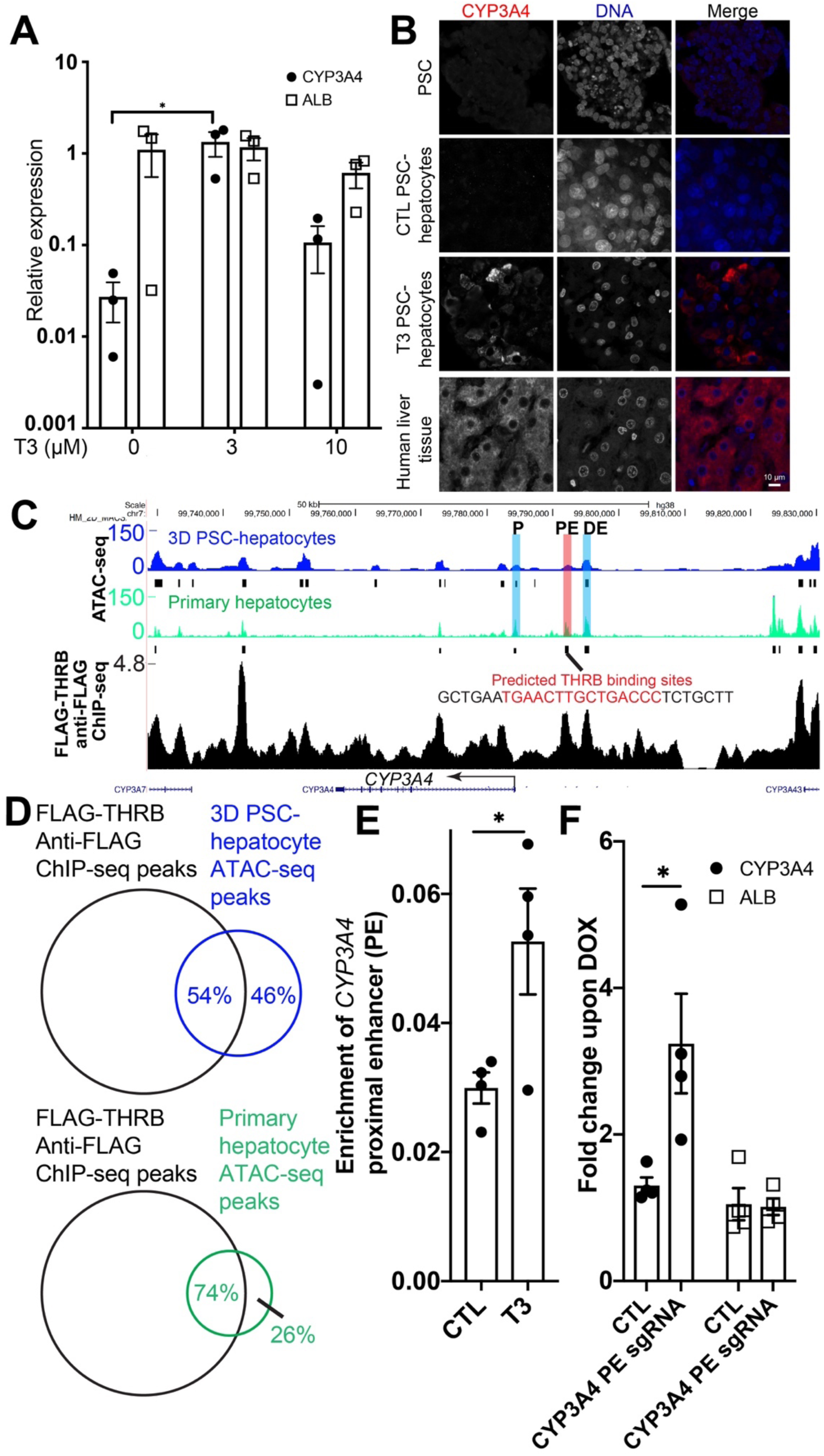
**Thyroid hormone depended upregulation of *CYP3A4* transcription is mediated by binding of HRB to the proximal enhancer of *CYP3A4*.** A) Effects of T3 on *CYP3A4* and *ALB* expression in PSC-hepatocytes differentiated in 3D spheroids ultures. Plotted data are mean ± s.e.m., *n*=3. * denotes *P*<0.05, two-sided Student’s *t* test. B) Immunofluorescence with an anti-CYP3A4 antibody (red) on undifferentiated PSCs, control 3D PSC-epatocytes, T3 treated 3D PSC-hepatocytes, and human liver tissue. C) ATAC-seq track of the *CYP3A4* locus was overlayed with anti-FLAG ChIP-seq data (GEO: SM2534017) from THRB-FLAG HepG2 cells (HepG2 cells with FLAG tagging to the endogenous *THRB* ene). The promoter and two 5’ enhancer elements of *CYP3A4* were denoted as P, PE (proximal enhancer), nd DE (distal enhancer). D) Overlapping of THRB ChIP-seq and ATAC-seq peaks from 3D PSC-hepatocytes (top panel) and primary epatocytes (bottom panel). The percentage of ATAC-seq peaks that overlapped with THRB ChIP-seq eaks were labelled. E) THRB-FLAB HepG2 cells were cultured with T3 or not (CTL), followed by cut-and-run experiment with an nti-FLAG antibody. The isolated DNA from MNase-proteinA treated cells were purified and subjected to q-CR analysis with primers spanning *CYP3A4* proximal enhancer (PE in c). Plotted data are mean ± s.e.m., =4. * denotes *P*<0.05, two-sided Student’s *t* test. F) DOX-inducible CRISPR activation H1 human PSCs were transduced with control sg lentivirus (CTL) or entivirus encoding sg RNA targeting *CYP3A4* proximal enhancer (PE). DOX was added to differentiated epatocyte-like cells for about 2 days, and RNA samples from cells were subjected to q-RT-PCR analysis ith *CYP3A4* primers (closed circles) and *ALB* primers (open squares). Plotted data are mean ± s.e.m., *n*=4. denotes *P*<0.05, two-sided Student’s *t* test. he scale bar in b: 10 µm.

We used THRB-FLAG HepG2 cells (Partridge et al., 2020) to further dissect the mechanisms of *CYP3A4* regulation by T3. Culturing of THRB-FLAG HepG2 cells in T3 containing medium resulted in increased CYP3A4 activity and *CYP3A4* expression (**Figure S4A-B**). When THRB-FLAG HepG2 cells were cultured in William’s E medium with different concentrations of T3, we found that 3 nM T3 mostly efficiently upregulated *CYP3A4* enzyme activity, and this was abolished with thyroid hormone receptor antagonist (1-850) (**Figure S4C-D**). Furthermore, endoribonuclease prep siRNA (esiRNA) knockdown of *THRB* showed that THRB was required for CYP3A4 upregulation by T3 (**Figure S4E**). To test THRB directly affects CYP3A4 activity by interacting with the proximal enhancer, we generated proximal enhancer deletion clones by CRISPR-mediated deletion (**Figure S4I-J**) and found that the upregulation of *CYP3A4* by T3 was reduced in the *CYP3A4* proximal enhancer deletion cells (**Figure S4L**). Additionally, we performed Cut-and-qPCR in THRB-FLAG HepG2 cells with anti-FLAG antibody with or without T3 using primers for the *CYP3A4* proximal enhancer. T3 increased the binding of THRB to the *CYP3A4* proximal enhancer compared to control (**Figure 3E**). These results suggest that THRB functions in regulating *CYP3A4* expression by binding to *CYP3A4* proximal enhancer in a T3-regulated manner.

To test if the mechanism applied to PSC-hepatocytes, we generated an inducible CRISPR activation PSC cell line (**Figure S5A-B**). Doxycycline (DOX) mediated induction of *PDX1*expression with a sgRNA targeting the *PDX1* promoter verified the inducible CIRPSR activation system (**Figure S5C-G**). We transduced the inducible CIRPSR activation PSCs with sgRNA targeting the *CYP3A4* proximal enhancer, and found that DOX added to hepatocyte spheroids generated from these cells increased *CYP3A4* expression but not another hepatocyte gene *ALB* (**Figure 3F**). These results suggest that thyroid hormone mediated THRB binding to the *CYP3A4* proximal enhancer to regulate *CYP3A4* expression.

### T3 enhances differentiation to more mature hepatocytes

In addition to adding T3, we removed oncostatin from the medium because oncostatin levels are downregulated upon birth (Kamiya et al., 2001), and we supplemented the media with antioxidant N-acetyl cysteine at the final stage of differentiation (**Figure S6A**). Quantitative-RT-PCR analysis of differentiation stages showed expected gene expression changes (**Figure S6B**). Immunofluorescence staining showed expression of HNF4A and ALB in most control cells and T3 treated cells (**Figure 4A**). Consistent with RNA expression levels, CYP3A4 protein levels increased, and AFP protein levels decreased upon T3 treatment (**Figure 4A**). KeyGenes analysis(Roost et al., 2015) of RNA-seq results showed that T3 treated hepatocytes were closer to primary hepatocytes than the control cells (**Figure 4B**), and T3 treatment increased expression of *CYP3A4* and *CYP2C9* (**Figure 4C-D**).

**Figure. 4.**
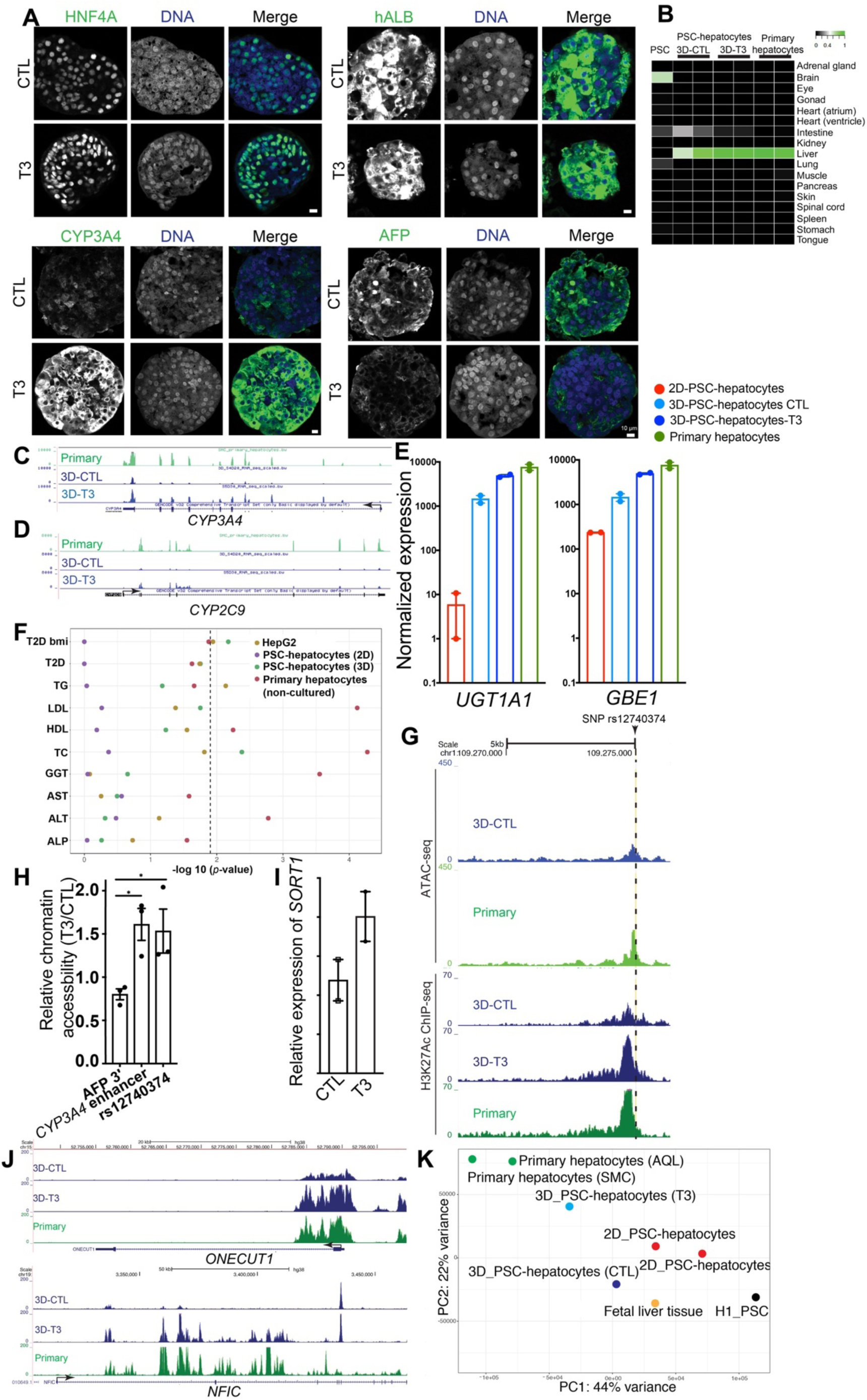
**Generation of PSC-hepatocytes with advanced maturity by modulating thyroid hormone ignaling.** A) Immunofluorescence staining of anti-HNF4A, human Albumin (ALB), CYP3A4, and AFP antibodies in ontrol PSC-hepatocytes (CTL) and PSC-hepatocytes treated with T3. B) KeyGenes analysis of gene expression comparison of 3D control PSC-hepatocytes, 3D PSC-epatocytes treated with T3, and primary hepatocytes. C-D) Genome browser gene tracks representing RNA-seq results of the *CYP3A4* (c) and *CYP2C9* (d) loci rom primary hepatocytes (Primary), control 3D PSC-hepatocytes (3D-CTL), and 3D PSC-hepatocytes reated with T3 (3D-T3). E) Expression of *UGT1A1* and *GBE1* in 2D-PSC-hepatocytes (red), control 3D-PSC-hepatocytes (light lue), T3 treated 3D-PSC-hepatocytes (dark blue), and primary hepatocytes (green). Plotted data are mean ± standard error, *n*=2. F) Comparison of enrichment of GWAS signal in open chromatin in 2D-PSC-hepatocytes, 3D-PSC-epatocytes, and primary hepatocytes versus open chromatin across all ENCODE and Roadmap pigenomic tissues for a set of liver-relevant diseases and traits. Dotted line indicates *P* <0.05 and colors enote the hepatocyte cell type tested. G) ATAC-seq tracks (top panel) and H3K27Ac ChIP-seq tracks (bottom panel) of the region surrounding a ommon non-encoding SNP rs12740374 that regulates *SORT1* expression in hepatocytes and plasma LDL-levels. H) Relative chromatin accessibility of *AFP* 3’ chromatin accessible peak, *CYP3A4* proximal enhancer peak, s12740374 peak from ATAC-seq libraries were measured by normalization to an *ACTB* chromatin ccessible peak with using quantitative-PCR. The ratio of PSC-hepatocytes treated with T3 comparted to ontrol PSC-hepatocytes were plotted as mean ± s.e.m. (*n*=3). I) Upregulation of *SORT1* expression by T3 treatment. Data shown are mean ± standard error, *n*=2. J) Gene tracks representing H3K27K27 ChIP-seq data of the *ONECUT1* and *NFIC* loci of control PSC-epatocytes (3D-CTL), T3 treated PSC-hepatocytes (3D-T3) and primary hepatocytes (Primary). K) PCA analysis of super-enhancers from H1 PSC, fetal liver tissue, 2D PSC-hepatocytes, 3D PSC-epatocytes treated with T3 (T3), control 3D PSC-hepatocytes (CTL), and primary hepatocytes (AQL and MC). Scale bars in a: 10 µm.

Further analysis of liver metabolic genes and disease relevant genes indicate that the PSC-hepatocytes generated with the optimized differentiation system were more mature. The drug metabolism gene *UGT1A1* and glycogen metabolism gene *GBE1*were upregulated in 3D-PSC-hepatocytes, and T3 treatment of 3D-PSC-hepatocytes further increased their expression (**Figure 4E**). Additionally, expression of glucose production regulator gene *G6PC*, clotting factors *F5*, and complement gene *C5* in 3D-PSC-hepatocytes were expressed at levels comparable to primary hepatocytes (**Figure S6C**). Furthermore, multiple genes encoding enzymes in the urea cycle (including the most commonly mutated gene in urea cycle disorder *OTC* and other genes encoding enzymes for the pathway: *NAGS*, *ASS1*, *ARG1*, and *ORNT1/SLC25A15*) were highly expressed in the 3D PSC-hepatocytes (**Figure S6D**), and urea levels were higher in 3D PSC-hepatocytes than 2D PSC-hepatocytes (**Figure S6E**). Moreover, the 3D PSC-hepatocytes increased expression of *INSR* and decreased expression of *IGF1R*, suggesting the 3D cells are better than 2D PSC-hepatocytes in studying metabolic effects of insulin signaling (**Figure S6F**).

In addition to protein-coding genes, we examined the enrichment of single nucleotide polymorphisms (SNPs, including non-coding SNPs) that are linked to liver-relevant diseases and traits in accessible chromatin regions of 2D-PSC-hepatocytes, 3D-PSC-hepatocytes, and primary hepatocytes. We applied stratified LD score regression (Finucane et al., 2018) to examine the summary statistics from 10 GWAS studies – ALP, ALT, AST, and GGT (Chambers et al., 2011); HDL, LDL, TC and TG (Willer et al., 2013); T2D and T2D stratified by body mass index (BMI)(Mahajan et al., 2018) and found that the enrichment of disease-associated SNPs in 3D PSC-hepatocytes was higher than those in 2D hepatocytes differentiation (**Figure 4F**). For T2D and BMI, the levels of enrichments in 3D PSC-hepatocytes and in primary hepatocytes were similar (**Figure 4F**), suggesting the 3D-PSC-hepatocytes could be used to study the disease-associated SNPs and the underlying mechanisms.

Furthermore, a common non-encoding SNP rs12740374 that regulated *SORT1* expression in hepatocytes by creating a C/EBP binding sites (Musunuru et al., 2010) and modulated plasma LDL-C levels was in an active ATAC-peak with H3K27Ac signals in 3D PSC-hepatocytes, similar to primary hepatocytes (**Figure 4G**). T3 treatment increased the expression and chromatin accessibility of *SORT1* (**Figure 4H-I**). Similarly, chromatin accessibility was increased for *CYP3A4* proximal enhancer, but reduced for *AFP*3’ region (**Figure 4H**). Additionally, T3 treatment improved H3K27Ac signals around rs12740374 (**Figure 4G**), and increased H3K27Ac signals of the *ONECUT1* and *NFIC* loci (**Figure 4J**). Analysis of super-enhancers also showed that T3 treated 3D-PSC-hepatocytes were more similar to primary hepatocytes than control 3D-PSC-hepatocytes (**Figure 4K**). These changes associated with hepatocytes maturation, together with extensive overlapping of THRB ChIP-seq peaks and ATAC-seq peaks (**Figure 3D**) indicate that the supplementing T3 could improve overall maturation status of PSC-hepatocytes.

To start examining the mechanisms by which THRB mediated the epigenetic changes, we performed immunoprecipitation (IP) of THRB in THRB-FLAG HepG2 cells with anti-FLAG antibody, followed by mass-spectrometry to identify THRB-binding proteins (**Figure 5A**). Consistent with a previous study showing interaction between THRB and heterogeneous nuclear ribonuclearprotein hnRNP H1 (Uren et al., 2016), multiple hnRNPs were identified in the immunoprecipitation complex (**Figure 5B**), in addition to several transcription initiation factor TAF proteins. Furthermore, PBRM1/BAF180 and ARID2/BAF200, two defining components of the ATP-dependent chromatin remodeling complex pBAF (polybromo-associated Brg/Brahma-associated factors (BAF)) (Kadoch and Crabtree, 2015) were identified to be present in the immunoprecipitation complex. To verify the mass-spectrometry results, we performed IP-Western experiments, and confirmed that PBRM1 and ARID2 were co-immunoprecipitated with THRB (**Figure 5C**). Furthermore, ARID2, loss-of-function, which has been frequently identified in hepatocellular carcinoma (Li et al., 2011), showed increased association to the IP-THRB complex in a T3 regulated manner (**Figure 5C**). These biochemical interactions between THRB and pBAF components, together with the effects of T3 on chromatin accessibility (**Figure 4H**), suggest that the pBAF complex could be involved in THRB mediated regulation of chromatin accessibility.

**Figure 5.**
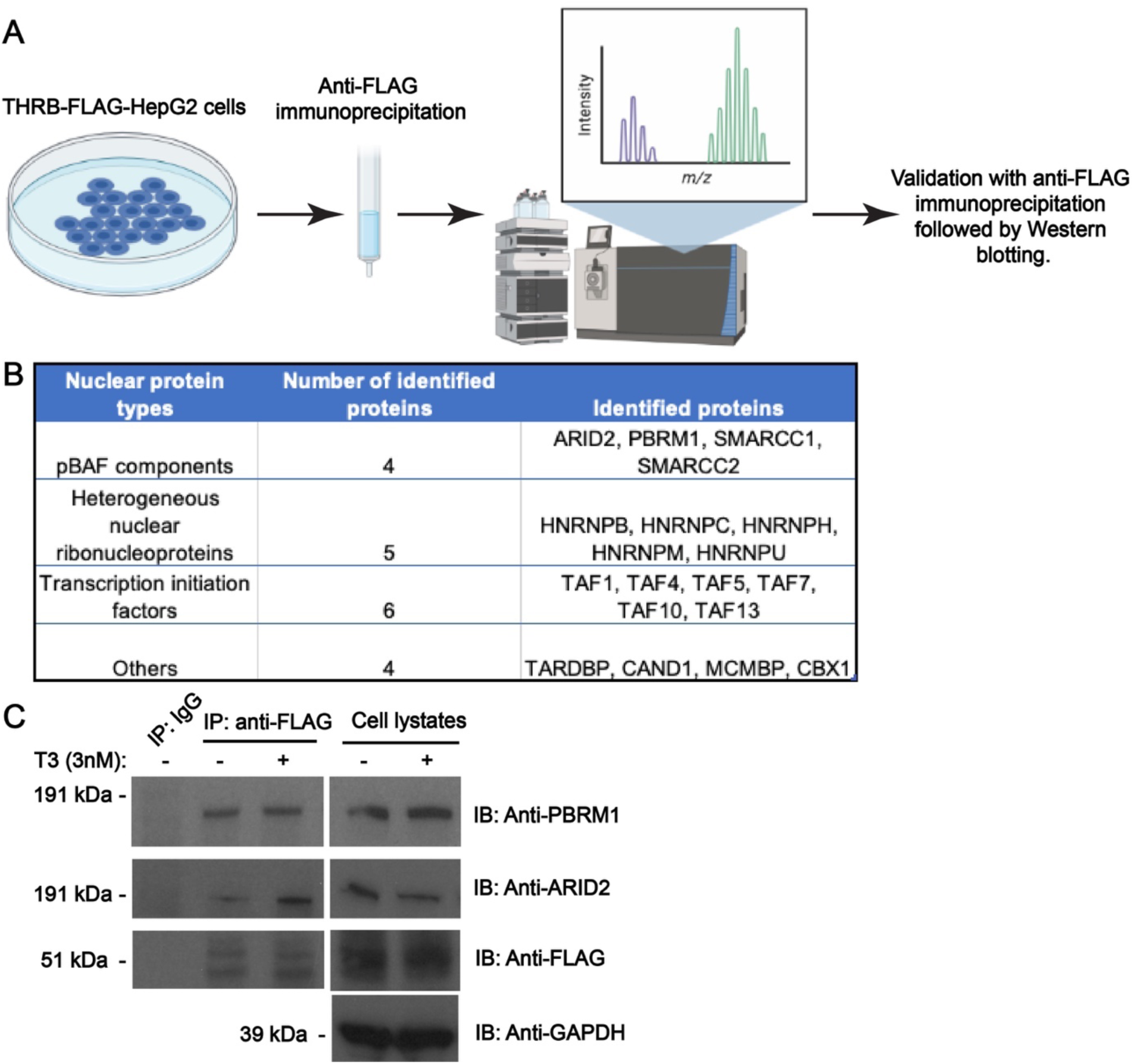
**Interactions between THRB and pBAF components in THRB-FLAG HepG2 cells.** (A) Experimental design to identify THRB-binding proteins by immunoprecipitation with anti-FLAG antibody in THRB-FLAG HepG2 cells followed by mass-spectrometry and IP-Western blotting analysis. (B) Identification of high confidence nuclear proteins co-immunoprecipitated with THRB-FLAB (*P*<0.01). (C) Control THRB-FLAG HepG2 cells or cells treated with 3 nM T3 for 1 day were lysed and subjected to anti-FLAG immunoprecipitation and Western blotting with pBAF components PBRM1 and ARID2.

We also tested the 3D spheroids differentiation for additional human PSC cell lines (**Figure S7A-C**). Gene expression analysis on 3D-PSC-hepatocytes generated from a tdTomato-expressing iPS cells derived from a female Niemann-Pick disease type C (NPC) patient (Maetzel et al., 2014), a GFP-expressing male ES cell line HUES8-GFP (Ma et al., 2018), and a female ES cell line WIBR3 (Lengner et al., 2010) showed reduced fetal hepatocyte gene *AFP* when 3D-PSC-hepatocyte were treated with T3 compared to control (**Figure S7D-E**). In contrast, expression of mature hepatocyte gene *CYP3A4* and *NFIC* were upregulated under this condition (**Figure S7E**) in the hPSCs. These results from different PSCs from different sources demonstrated that this approach supported differentiation of multiple hPSC lines including patients’ iPS cells to more mature hepatocytes.

### Engraftment of hPSC-hepatocytes to the undamaged liver of immunocompromised mice

Since hepatocytes maturity is related to the capacity to engraft into liver upon transplantation (Hu et al., 2018), we tested the *in vivo* function of the PSC-differentiated hepatocytes by transplanting the cells into immunocompromised mice. Transplantation of human hepatocytes or human cells derived hepatocytes have been performed in immunocompromised mice with liver injuries to facilitate engraftment of human cells(Azuma et al., 2007; Carpentier et al., 2014; Hu et al., 2018; Michailidis et al., 2020; Xie et al., 2019). Here, we tested if PSC-differentiated hepatocytes in 3D could be engrafted in undamaged mouse liver by injecting dissociated cells into the spleen of NOD.Cg-Prkdcscid Il2rgtm1Wjl/SzJ (NSG) mice (**Figure 6A**). H&E histology staining showed that liver histology of transplanted mice both 1 month post transplantation and 6 months post transplantation were similar to control mice (**Figure 6A**). However, anti-human-Albumin antibody IHC detected engrafted human cells with hepatocyte morphology integrated to the mouse liver parenchyma (**Figure 6B**). Furthermore, immunofluorescence staining showed that 1 month after transplantation, PSC-hepatocytes expressing human ALB and CYP3A4 could be found in liver of transplanted mice (**Figure 6C**). At 6 months post transplantation, human albumin expressing hepatocytes could be detected to integrate with mouse liver parenchyma (**Figure 6D**). We extracted DNA from tissue slides and performed q-PCR analysis and detected human mitochondrial DNA (**Figure 6E**), consistent with the presence of human hepatocytes in the transplanted mouse liver. These data show that that 3D PSC differentiated hepatocytes can engraft into undamaged liver of immunocompromised mice and integrate into the liver parenchyma without disrupting the normal histology of the liver.

**Figure 6.**
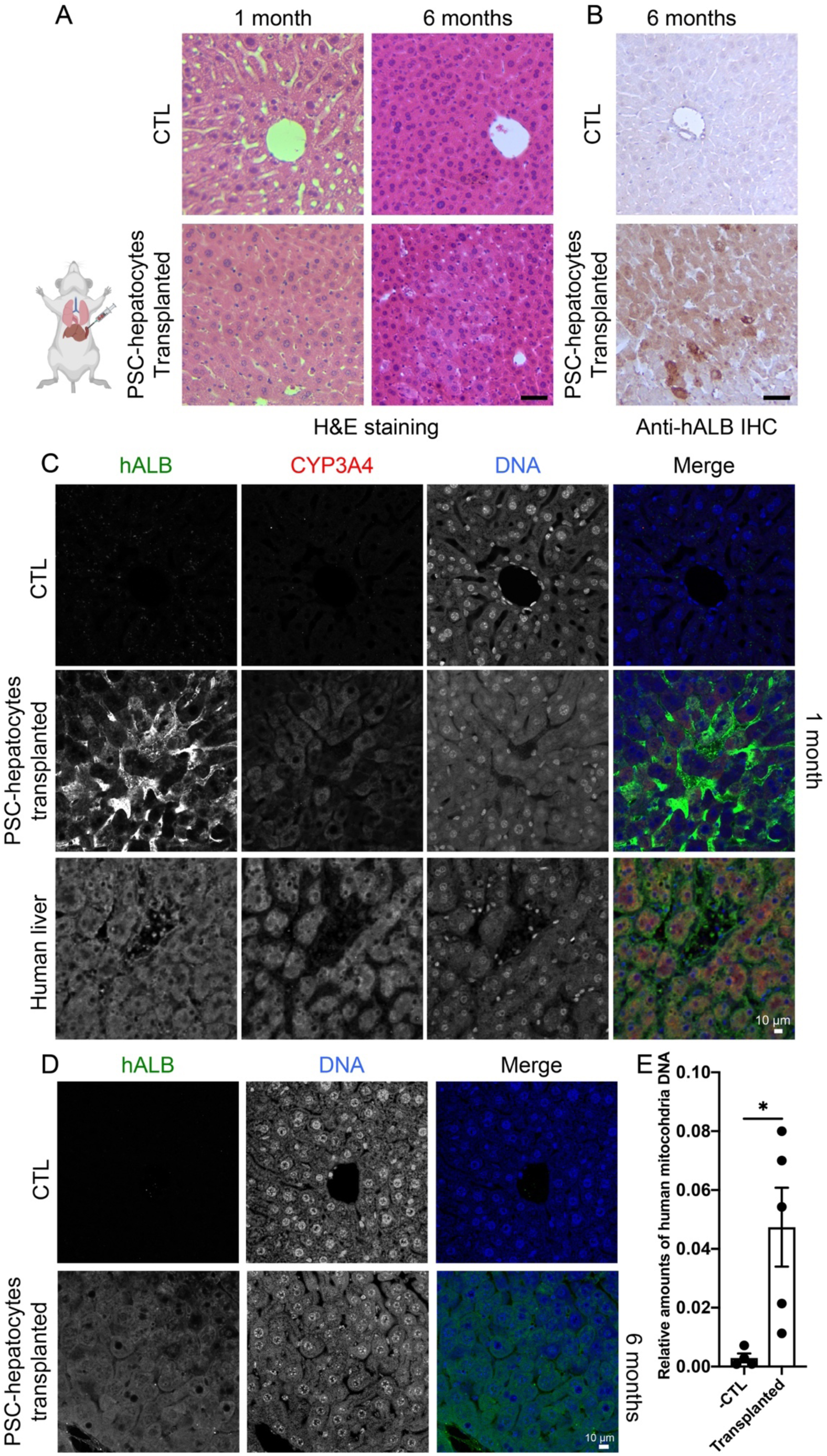
**Engraftment of 3D PSC-hepatocytes into undamaged mouse liver.** (A) H&E staining of liver from control mice (top panels) or from mice with splenic transplantation of dissociated human 3D PSC-hepatocytes (bottom panels) at 1 month post transplantation (left panel) or 6 months post transplantation (right panels). (B) Anti-human albumin (hALB) immunohistochemistry of liver from control mice (top panels) or from transplanted mice (bottom panels) at 6 months post transplantation. (C) Immunofluorescence with an anti-human albumin (hAlbumin) antibody (green) and an anti-CYP3A4 antibody (red) on control mouse liver (top panel), liver from transplanted mice (middle panel) 1 month post transplantation, or human liver (bottom panel). (D) Immunofluorescence with an anti-human albumin antibody staining (green) of control mouse liver (top panel), or liver from transplanted mice (bottom panel) 6 months post transplantation. (E) Quantification of human mitochondrial DNA from DNA samples isolated from slices from control mice or transplanted mice. Plotted data are mean ± s.e.m., *n*=4 for control, *n*=5 for transplanted mice. * denotes *P*<0.05, two-sided Student’s *t* test. Scale bars in (A) and (B): 50 µm. Scale bars in (C) and (D): 10 µm.

## Discussion

In this study, we compared genome-wide maps of gene expression and cis-regulatory elements of PSC-hepatocytes with that of primary non-cultured hepatocytes. Modulating thyroid hormone signaling, contributed to the generation of more mature human hepatocytes. THRB regulated *CYP3A4* expression by binding to enhancer elements through ligand-regulated binding and activation. The PSC-hepatocytes not only expressed critical hepatocyte genes, but also showed activation of disease associated non-coding SNPs. Upon transplantation to uninjured liver of immunocompromised mice, these PSC-hepatocytes integrated into mouse liver parenchyma without disrupting normal liver histology for up to 6 months after transplantation. These data suggest that PSC-hepatocytes could be used to study expression and regulatory mechanism of hepatocyte genes implicated in hepatocyte development and liver disease. In addition, these PSC-derived hepatocytes may be useful for cell replacement therapies.

Our results are consistent with the critical role of thyroid hormone signaling for liver physiology. Hypothyroidism has been connected with hyperlipidemia(Willard et al., 2014), and T3 treatment was reported to improve liver dysfunction associated with congenital hypothyroidism(Mantri et al., 2016). Improved H3K27Ac signals upon T3 treatment may reflect recruitment of histone acetylase p300/CBP to nuclear hormone receptors (Ogryzko et al., 1996). It is possible that thyroid hormone receptors recruit pBAF to regulate chromatin accessibility through nucleosome repositioning or ejection (Clapier et al., 2017), similar to vitamin D receptors that switch pBAF complex for BAF complex in pancreatic β cells (Wei et al., 2018). The thyroid hormone signaling is regulated at multiple levels including expression of thyroid hormone transporter proteins and T3 metabolizing enzymes such as iodothyronine deiodinases. Although the effects of thyroid hormones could be context depended, our results suggested the critical functions of thyroid hormone signaling in promoting maturation of human PSC-differentiated hepatocytes. Based on the conserved functions of thyroid hormones in regulating organ development (Brent, 2012), and the surge of thyroid hormone levels at and shortly after birth, T3 has been tested empirically to improve maturation of iPS differentiated hepatocytes in 2D culture (Bogacheva et al., 2021). Our unbiased transcriptome and epigenome study using 2D and 3D differentiations identified that thyroid hormone signaling accounted for a significant portion of the immaturity phenotype of hepatocytes differentiated from hPSC *in vitro*. Furthermore, we provided evidence suggesting that THRB could bind the chromatin remodeling complex pBAF, a potential mechanism by which THRB could mediate chromatin accessibility changes of gene regulatory elements. The environmental factors-nuclear receptors axis could regulate maturation of additional cell types differentiated from human PSCs.

Future studies using 3D chromosome conformation capture methods and single cell genomics and epigenomics approaches on the PSC-hepatocytes could further provide insights into the developmental and maturation processes of human hepatocytes. Furthermore, scaling up PSC-hepatocytes differentiation with bioreactors and potentially microencapsulation, modulating additional nuclear receptors signaling pathways to further improve functional maturation of PSC-hepatocytes, and expansion of the PSC-hepatocytes *in vitro* and *in vivo* in immunocompromised mice with liver damages could be future studies.

## Methods

### Primary human liver tissue sections

Formalin-fixed paraffin-embedded (FFPE) slides (5-micron sections) from a normal adult human liver were obtained from Biomax (HuFPT074 SB, male, 45 years old). The antigen retrieval and immunofluorescence staining procedures were performed with a modified protocol (Li et al., 2014). Briefly, after deparaffinization and rehydration, the FFPE slides were subjected to antigen retrieval with citrate buffer (10 mM, pH 6.0). Then the tissue slides were blocked in PBST (PBS with 0.1% Tween-20) and 5% donkey serum (Jackson Laboratories 017-000-121) for 2 hours at room temperature, followed by primary antibodies (1:100 mouse-anti-CYP3A4 antibody Life Technologies MA5-17064, 1:100 goat-anti-hALB antibody, Bethyl Laboratories A80-229A, diluted in PBST with 5% donkey serum) incubation at 4 °C for overnight. After four times washing with PBST for 10 minutes each at room temperature, secondary antibodies (Alexa Fluor 647 conjugated donkey anti-goat IgG secondary antibody Life Technologies A21447 and Alexa Fluor 594 conjugated donkey anti-mouse IgG secondary antibody Life Technologies A21203, 1:400 diluted in PBST with 5% donkey serum) were added for incubation at room temperature for 1 hour. After four times washing with PBST for 10 minutes each, DAPI staining was for 10 minutes at room temperature followed by rinsing with PBST for 3 times. After mounting with Fluoromount-G mounting medium (Electron Microscopy Sciences 1798425), the slides were imaged with a Zeiss LSM 710 confocal microscope.

### Primary human hepatocytes

Frozen primary human hepatocytes from healthy donors were purchased from BioIvt (Catalog number M00995-P, lot: SMC and AQL). After thawing in a 37 °C water bath, the hepatocytes were washed with Cryopreserved Hepatocyte Recovery Medium (CHRM, Thermo Fisher Scientific, CM7000). RNA from about 0.5 million cells were purified with RNeasy Plus Micro (QIAGEN). Around 50,000 cells were washed with PBS and subjected to tagmentation and ATAC-seq described below. About 5-10 million cells were crosslinked with 1% formaldehyde for ChIP-seq described below.

### Culture of human PSCs

Matrigel coated tissue culture plates were used for feeder-free culture of human PSCs. For coating plates, Matrigel (Corning) was 1:100 diluted in cold DMEM-F12 medium, and 2 ml of diluted Matrigel solution was added to one well of a 6-well plate well, and coat for overnight at room temperature before pre-warming in a 37 °C incubator for 1 hour prior to plating cells. Human PSCs H1-OCT4-GFP(Zwaka and Thomson, 2003), H1-OCT4-GFP AAVS1:tdT(Ma et al., 2018), HUES8-GFP(Ma et al., 2018), and INS:tdT(Ma et al., 2018) were cultured in mTeSR medium (STEMCELL Technologies) in Matrigel coated 6-well plates, and passaged very 3-4 days with a 1:4-6 passage ratio using Versene solution (Life Technologies 15040066).

WIBR3(Lengner et al., 2010) and Niemann-Pick disease type C (NPC1) iPS (clone 16-13(Maetzel et al., 2014)) were initially cultured on mitomycin C inactivated mouse embryonic fibroblasts (MEFs) feeder cells using DMEM-F12 (Life Technologies cat# 11330-057) supplemented with 15% FBS (Hyclone SH30396.03), 5% KSR (Life Technologies cat# 10828-028), 4 ng/ml FGF (Life Technologies cat# PHG0261), 0.1 mM 2-mercaptoethanol (Life Technologies cat #21985-023), 1X L-glutamine (Life Technologies cat# 25030-081), 1X MEM-NEAA (Life Technologies cat# 11140-050), 1X penicillin/streptomycin (Life Technologies cat# 15140-122). Cells were passaged with 1 mg/ml collagenase IV (Life Technologies cat# 17104019) every 4-6 days on inactivated MEF feeders. All human PSCs were routinely tested negative for mycoplasma.

### Genetic modification of human PSCs

NPC-iPS cells (clone 16-13(Maetzel et al., 2014)) were targeted with AAVS1-tdTomato (Addgene 159275) as described before(Ma et al., 2018). Briefly, NPC-iPS cells were treated with 10 µM Rho kinase inhibitor Y-27632 overnight before dissociated to single cells with Accutase (Stem Cell Technology 07922). About 10 million cells were electroporated with 10 μg of AAVS1 zinc finger nuclease-encoding plasmid and 30 μg of AAVS1-tdTomato with one electroporation of 250 V, 500 μF using a Gene Pulser Xcell System (Bio-Rad), and plated on DR4 MEFs in PSC medium with 10 µM Y-27632 during the first day. Puromycin (0.5 μg/mL) selection was added 3–4 days after electroporation to select resistant cells. The tdTomato-expressing clones were manually passed to inactivated MEF feeders plated in 12-well plates in regular PSC medium without puromycin.

DOX-inducible CRISPR-activation cells were generated based on the INS:tdT reporter H1 cells(Ma et al., 2020) first by removing the floxed puromycin resistant cassette by transient expression of Cre. About 3 million Y-27632 pretreated H1-INS:tdT PSC cells were electroporated with 5 µg CAGGS:Cre-tdT under electroporation condition stated above and plated in mTeSR medium supplemented with 10 µM Y-27632 in Matrigel coated plates. One day after electroporation, tdTomato-expressing cells were isolated by fluorescence activated cell sorting (FACS) using a FACSAria cell sorter (BD Biosciences) and plated sparsely on Matrigel coated plates in mTeSR and 10 µM Y-27632. Individual clones were plated in 12-well plates and puromycin-sensitive clones were expanded and targeted with an AAVS1-DOX inducible CRISPR activation plasmid. Puromycin (0.5 µg/mL) resistance clones were isolated after targeting. Isolated clones were characterized by Southern blotting using the *AAVS1* probes as described before(Ma et al., 2020), and properly targeted cells were expanded.

HEK293T cells were cultured in HEK293 medium (DMEM supplemented with 10% FBS, 1X L-glutamine, 1X MEM-NEAA, 1X penicillin/streptomycin), and passaged with trypsin-EDTA. Lentiviruses delivering sgRNA were packed by transfecting about 70% confluent HEK293T cells cultured in T75 flasks with 5 µg pCMV-VSVG (Addgene 8454), 10 µg pCMV-dR8.2 dvpr (Addgene 8455), and 10 µg sgRNA plasmid using X-tremeGENE™ 9 DNA transfection reagent (Roche 06365809001). One day after transfection, medium was changed to HEK293T culture medium. Media were changed daily, and the conditioned media were collected for 2-3 days, filtered through 0.45 µM filters and ultracentrifuged (23000 rpm at 4 °C for 1 hour 45 minutes) with a Beckman Optima XE-90 ultracentrifuge. Centrifuged pellets were resuspended in 500 µL PBS overnight at 4 °C. For lentiviral transduction, DOX-inducible CRISPR activation H1 cells were pretreated with 10 µM Y-27632 overnight, then single cell dissociated with Accutase, and plated in Matrigel-coated 6-well plates in 2 ml mTeSR medium supplemented with10 µM Y-27632, and 20 µL CRISPR activation sgRNA virus stock. After 1 day, the medium was changed to mTeSR medium. Zeocin selection (0.2 mg/mL) was started after another 2 days for about 3-5 days. Zeocin resistant cells were passaged 1:4 in mTeSR medium with zeocin for about 1 more week. The cells were expanded in mTeSR for differentiation.

### Differentiation of human pluripotent stem cells (PSCs) to hepatocytes-like cells

WIBR3 cells and NPC-iPS-tdT cells were adapted to feeder culture with mTeSR1 medium by manual passaging. Cells were differentiated to hepatocyte-like cells after 3-6 passages in mTeSR medium.

All feeder-free human PSCs were thawed in mTeSR1 medium, and cultured as described before. Cells were typically differentiated within 3 passages after thawing.

For 2D differentiation of H1-OCT4-GFP cells, less than 60% confluent PSCs were treated with mTeSR1 medium supplemented with 10 μM Y-27632 for overnight or for 4-6 hours and the cells were dissociated into single cells with Accutase for about 5 min at 37°C by gentle pipetting in mTeSR1 medium. After cell counting with Countess (Invitrogen), about 1.4-1.6 million live cells were plated to each well of a 6-well Matrigel-coated plates in mTeSR medium supplemented with 10 μM Y-27632 and distributed evenly. After 1 day when plated cells were about 80-90% confluent, differentiation was initiated by removing mTeSR medium, washing cells with DMEM/F12, and followed by daily media changes (stages and days are referred by S and D, respectively):

Day 1 (S1D1): MCDB131 medium with 10mM Glucose,1.5 g/L NaHCO3, 0.5 % FAF-BSA, 1x Glutamax, 1x penicillin/streptomycin, 100 ng/ml activin, and 3 μM Chir99021.

Day 2 (S1D2): MCDB131 medium with 10mM Glucose,1.5 g/L NaHCO3, 0.5 % FAF-BSA, 1x Glutamax, 1x penicillin/streptomycin, 100 ng/ml activin, and 0.3 μM Chir99021.

Day 3 (S1D3): MCDB131 medium with 10mM Glucose,1.5 g/L NaHCO3, 0.5 % FAF-BSA, 1x Glutamax, 1x penicillin/streptomycin, and 100 ng/ml activin.

Day 4-8 (S2D1-S2D5): Cells were washed with PBS on S2D1 before changing to RPMI-1640 medium with 1x glutamax, 1x NEAA, 1x penicillin/streptomycin, 1x B27, 10 ng/ml FGF2, and 20 ng/ml BMP4.

Day 9-13 (S3D1-S3D5): RPMI-1640 medium with 1x glutamax, 1x NEAA, 1x penicillin/streptomycin, 1x B27, and 20 ng/ml HGF.

Day 14-18 (S4D1-S4D5): William E medium supplemented with 1x glutamax, HCMTM SingleQuots Kit (no EGF was added), trace element A (500 mL), trace element B (500 ml), 100 nM dexamethasone, 5.35 mg/mL linoleic acid, 20 ng/ml HGF, and 20 ng/ml Oncostatin M.

For 3D spheroid differentiation of H1-OCT4-GFP cells, cells plating and media changes for the first 3 days were the same as described above. After S1D3, the following media changes were used:

Day 3-5 (S2D1-D2D2): S1D3 cells were washed with PBS before changing to S2 medium: MCDB131 with 0.5% FAF-BSA, 1.5 g/L NaHCO3, 10 mM glucose, 1x penicillin/streptomycin, 1x glutamax, 0.25 mM vitamin C, and 50 ng/ml KGF.

Day 6 (S3D1): RPMI-1640 medium with 1x glutamax, 1x NEAA, 1x penicillin/streptomycin, 1x B27 (1:50), 50 µg/ml heparin, 10 ng/ml FGF2, and 20 ng/ml BMP4.

Day 7-10 (S3D2-S3D5): After about 1 day incubation in S3D1, 10 μM Y-27632 was added to the medium for about 4-6 hours. Then cells were washed with PBS, and dissociated with Accutase for about 5 min at 37 °C. Then cells were washed with RPMI-1640 medium with 1x glutamax, 1x NEAA, 1x penicillin/streptomycin, 1x B27, and 1% BSA-fragment V. Approximately 2 million live cells were plated in 4 ml S3 medium (RPMI-1640 medium with 1x glutamax, 1x NEAA, 1x penicillin/streptomycin, 1x B27, 10 ng/ml FGF2, and 20 ng/ml BMP4 supplemented with 10 μM Y-27632 in each well of 6-well AggreWell 400 plates (Stem Cell Technology, 34415). Next day the formed cell spheroids clusters were washed with RPMI-1640 medium with 1x glutamax, 1x NEAA, 1x penicillin/streptomycin, 1x B27, and moved to 6-well ultralow adherent culture plates placed on an orbital shaker set at 95-100 rpm for 4-5 more days in S3 medium with daily media change. The rest of the differentiation were carried out in ultralow attachment plates placed on an orbital shaker with the same setting. During this and later steps, cystic spheroids and aggregated spheroids were removed daily.

S4 (about 2-3 weeks): William E medium supplemented with 1x glutamax, HCMTM SingleQuots Kit (no EGF was added, and 5 µL insulin was used for 500 mL medium), trace element A (500 μL), trace element B (500 μL), 100 nM dexamethasone, 100 nM gamma secretase inhibitor XX, 20 μM forskolin, 1 μM TGFβ inhibitor A8301, 5.35 μg/mL linoleic acid, 20 ng/ml HGF, and 20 ng/ml Oncostatin M. N-acetyl cysteine (1 mM) was added to the later part of S4 as needed to maintain culture health. Media were changed every 2 days.

S5 (about 1-2 weeks): William E medium supplemented with 1x glutamax, HCMTM SingleQuots Kit (no EGF was added, and 5 µL insulin was used), trace element A (500 μL), trace element B (500 μl), 100 nM dexamethasone, 100 nM gamma secretase inhibitor XX, 20 μM forskolin, 1 μM TGFβ inhibitor, 5.35 μg/mL linoleic acid, 20 ng/ml HGF, 3 μM T3, and 1 mM N-acetylcysteine. Media were changed every 2 days.

Cell growth state and density optimizations are needed for proper differentiation. Representative Q-RT-PCR results during the differentiation are show in Extended Data Fig. 9b.

For DOX-induction experiments, DOX-inducible CRISPR activation H1 cells were differentiated as above, and DOX (final concentration: 2 mg/ml) was added from S4D12 to S4D14. RNA samples were purified with RNeasy Plus Micro Kits (Qiagen) for q-PCR analysis (described later) using *ACTB*, *CYP3A4*, *ALB* and *cas9* primers.

For differentiating additional PSCs, the following seeding density and KGF concentrations for S2 were used: HUES8-GFP (1.4-1.6 million cells per 6-well plate well, 10 ng/ml KGF for S2), NPC-iPS-tdT (1.8-2 million cells per 6-well plate well, 5 ng/ml KGF for S2), WIBR3 (1.3-1.5 million cells per 6-well plate well, 10 ng/ml KGF for S2).

### Culture of THRB-FLAG HepG2 cells and deletion of *CYP3A4* proximal enhancer

THRB-FLAG HepG2 cells were maintained in DMEM with 10% FBS, 1x penicillin/streptomycin, 1x MEM-NEAA, 1% glutamax. Cells were passaged 1:4-1:6 about twice a week. THRB-FLAG HepG2 cells were routinely tested negative for mycoplasma.

To generate THRB-FLAG HepG2 cells with *CYP3A4* proximal enhancer deleted, 1 µg of 330-cherry-CYP3A4 enhancer-L-sgRNA, and 1 µg of 330-BFP-CYP3A4-enhancer-R-sgRNA were transfected with X-tremeGENE 9 DNA transfection reagent (Roche 6365809001). After 3-5 days, transfected cells were dissociated to single cells with Trypsin-EDTA, and cells with high expression of both cherry and BFP were isolated with fluorescence activated cell sorting with a FACSAria cell sorter (BD Biosciences). Clones grown from sorted cells were expanded and DNA samples from individual clones were extracted with Lucigen quick DNA buffer (68 °C for 15 min followed by 98 °C for 2 min), and genotyped by PCR with primers (F: GGTCCCCTTGGAACTTCATGC, R: CCTTCAACAACTAATAGCAGGG) with 2x KAPA HiFi HotStart ReadyMix (Roche) using the following PCR program (95 °C for 3 min; then 35 cycles of 98°C for 20 sec, 58°C for 15 sec, 72 °C for 30 sec; then 72 °C for 5 min). Control cells showed amplicon of about 0.5 kb, and clones with enhancer deletion showed amplicon of about 0.2 kb. PCR amplicons were sequenced by Sanger sequencing and clones with confirmed homozygous disruption of *CYP3A4* proximal enhancer were kept.

### Nuclear receptor ligands treatment and CYP3A4 activity measurements

For nuclear receptor ligands treatment, cells were cultured in William E medium supplemented with 1x glutamax, HCMTM SingleQuots Kit (no EGF was added, and 5 microliters of insulin was used for 500 ml medium), trace element A (500 µL), trace element B (500 µL), 5.35 µg/mL linoleic acid with different concentration of T3 (3nM-10µM) or retinoic acid (20 nM-1µM). For thyroid hormone antagonist experiments, a final concentration of 3 µM antagonist (1-850) was used with 3 nM T3.

For measuring CYP3A4 activity with CYP3A4 P450-Glo (Promega), IPA-luciferin was diluted 1:1000 in William E medium supplemented with 1x glutamax, HCMTM SingleQuots Kit (no EGF was added, and 5 microliters of insulin was used for 500 ml medium), trace element A (500 µL), trace element B (500 µL), 5.35 µg/mL linoleic acid and incubated in a 37 °C cell culture incubator. After 2 hours, 50 µL conditioned medium from each sample was mixed with 50 µL reconstituted luciferin detection reagent in a white opaque 96-well plate and incubated at room temperature for 20 min before data collection with a SpectraMax microplate reader (Molecular Devices).

### Transfection of THRB-FLAG-HepG2 cells with esiRNA

About 0.1-0.2 million THRB-FLAG HepG2 cells were plated in each well of a 24-well plate in 500 µL DEME medium with 10% FBS. The next day, control (SIC001, Millipore Sigma) or esiRNA targeting *THRB* (EHU083461, Millipore Sigma) were transfected using Lipofectamine RNAiMAX Transfection Reagent (Invitrogen) in 100 µL Opti-MEM medium with final concentration of 30 nM. After 2 days, media were removed and washed with PBS, and 500 µL William E base medium with 3 nM T3 or control were added. After 1 day, CYP3A4 luciferase experiments were performed as described above.

### Cut-and-qPCR experiments

About 70% confluent FLAG-THRB HepG2 cells cultured in 6-well plates were washed with PBS, and changed to William’s E base medium with or without 3 nM T3. After 1-2 days, cells were dissociated to single cells with trypsin, and 250,000 cells were used for each experiment based on the protocol(Skene et al., 2018). Specifically, buffers with 0.025% digitonin was used. Primary antibody (1 µg rabbit-anti-FLAG, Sigma F7425) or rabbit normal serum were incubated for 2 hours at 4 °C. After washing, 0.5 µL pA-MN (a gift generously provided by Dr. S Henikoff) was added to incubate at room temperature for 10 min.

Cutting experiments were performed at 0 °C ice-water bath with 50 µL buffer with CaCl_2_ for 30 min. Then 34 µL stop buffer were added and incubated at 37 °C for 15 minutes. DNA from collected supernatant were purified with ChIP DNA concentration and cleaning kit (Zymo D5205) and eluted in 20 µL elution buffer. For DNA input, 500,000 cells were used for genomic DNA purification with blood and tissue DNA kit (Qiagen 69504). Quantitative PCR experiments were performed with fast SYBR Green Master Mix (Thermo Fisher Scientific 4385618) with the *CYP3A4* proximal enhancer primers (F: CTTGCTGACCCTCTGCTTTCC; and R: CTCATGAGGCTGCTTGAACCG). Relative enrichment is calculated by normalization with input DNA.

### Immunoprecipitation, mass-spectrometry and Western blotting

THRB-FLAG-HepG2 cells were cultured in 10-cm cell culture plates in DMEM with 10% FBS, 1x penicillin/streptomycin, 1x MEM-NEAA, 1% glutamax. Cells, about 50-70% confluent, were washed with PBS, and 10 ml William’s E medium was added for 1-2 days. Then the cells were washed with PBS, and collected with 500 µL NP40 buffer (1% Nonidet P-40 (Igepal A-680), 50 mM Tris-HCl (pH 7.4), 150 mM NaCl, 2 mM EDTA, 50 mM NaF, 10 Glycerol 10 %, with freshly supplemented protease inhibitors (Roche 04693159001)). Cell lysates were collected, incubated at 4 °C for 30 min, passed through 1 ml syringe with 23G needle 10 times. Then spun at 14,000 rpm for 5 min. Then the pellets were ground for about 2 min, spun at 14,000 rpm for 5 min. The nuclear fraction and supernatant fraction (800 µL) were collected, and 8 µg rabbit-anti-FLAG antibody (Sigma F7425), or about 8 µg rabbit control serum were added to incubate overnight at 4 °C. The clear lysates were incubated with 100 µL protein G dynabeads (Thermo Fisher 10003D) for 2 hours at 4°C. Pellets were washed 4 times with NP40 buffer.

For mass-spectrometry, the dynabeads samples were washed and resuspended in a Tris/ Urea buffer, reduced with dithiothreitol, alkylated with iodoacetamide and digested with trypsin at 37 °C overnight. This solution was subjected to solid phase extraction to concentrate the peptides followed by injection onto a Thermo EASY-nLC 1200 HPLC equipped with an EASY-Spray ES900 3 um C18 analytical column 0.075 mm by 15 cm, (Thermo). Peptides were eluted using standard reverse-phase gradients. The effluent from the column was analyzed using a Thermo Exploris 480 mass spectrometer (nanospray configuration) operated in a data dependent manner for the 90 minutes. The resulting fragmentation spectra were correlated against the Refseq entries for homo sapiens using PEAKS Studio X+ (Bioinformatic Solutions) and to provide consensus reports for the identified proteins.

Western blotting was performed with a rabbit anti-PBRM1 antibody (Bethyl Laboratories A301-591A, 1:2000), a rabbit anti-ARID2 (Cell Signaling Technology 82342S 1:1000), or a mouse anti FLAG antibody (Sigma F1804 1:4000) based on previous methods (Wu et al., 2007).

### RNA purification and quantitative RT-PCR

PSC hepatocytes spheroids were washed with PBS, then lysed in RLT plus buffer from Qiagen RNeasy plus micro kit for about 15-20 min at room temperature followed by RNA purification. Reverse transcription was performed using by adding 1.5 µL q-cript to 6 µL RNA, and the reaction was carried out at 25 °C for 5 min, 42 °C for 30 min, 85 °C for 5 min. Quantitative PCR experiments were performed with SYBR Green Master Mix reagent with a QuantStudio 6 Flex Real-Time PCR System (Applied Biosystems). Relative quantifications were determined with the above primer pairs using delta delta threshold cycle methods with the following primers:

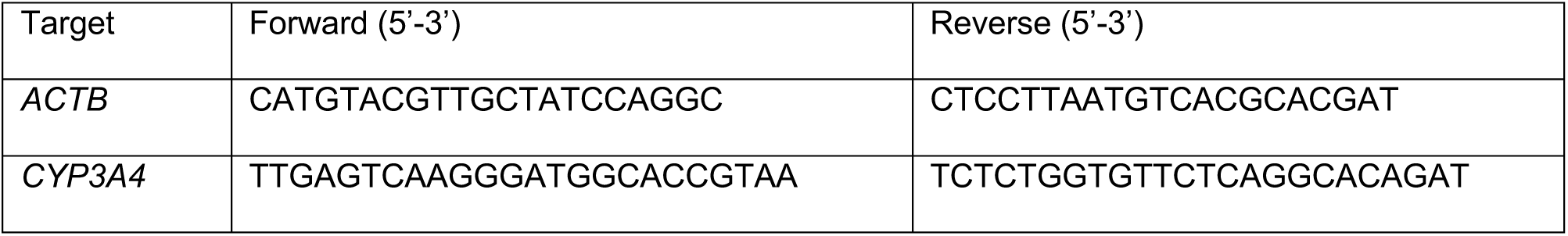
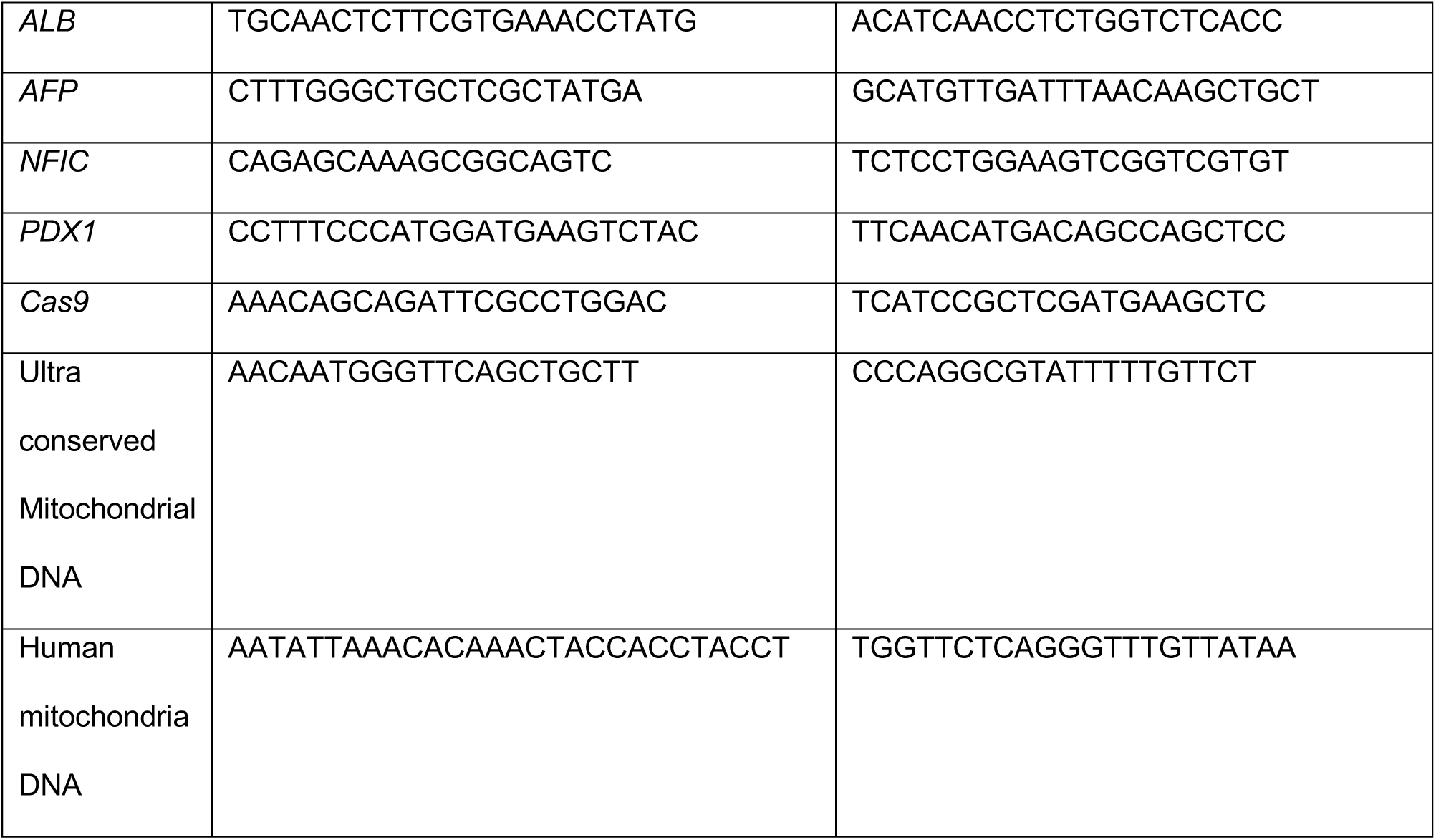

### RNA-seq and analysis

RNA samples from freshly thawed human primary hepatocytes (SMC and AQL) or from PSC-hepatocytes were prepared with RNeasy plus micro kit (Qiagen), and 10-20 ng total RNA samples were used for constructing stranded RNA libraries with Swift RNA-seq library preparation kit and sequenced with an Illumina HiSeq 2500 sequencer with 50 base single end reads.

Human fetal liver RNA-seq (GEO accession ID: GSM3067803) was processed with RNA-seq results generated in this study. Briefly, RNA-seq reads were aligned to human (GRCh38) genome using STAR(Dobin et al., 2013) V2.7.1a. Gene counts were obtained using featureCounts(Liao et al., 2014), and differentiation expression analyses were performed using DEseq2(Love et al., 2014). Enhanced volcano plots were generated in RStudio. Gene set enrichment analyses were performed using GSEA(Subramanian et al., 2005) 4.0.1, and KEGG pathway enrichment analyses were performed using g:profiler (https://biit.cs.ut.ee/gprofiler/). BAM files were converted to normalized Bigwig files using bamCoverage from deepTools(Ramirez et al., 2014) for visualization on a customized UCSC genome browser(Kent et al., 2002) session.

### Tn5 tagmentation for ATAC-seq

About 60-80 spheroids were washed with PBS, and dissociated with Accutase for about 20-25 min with gentle agitation in a 37 °C water bath. Then cells were gently pipetted, and washed with William’s E medium with 1% BSA-fragment V. After counting, 50,000 cells were centrifugate for 5 min at 500 RCF. Then the cells were washed with 50 µL cold PBC at 4°C for 500 RCF for 5 min. After removing PBS, 50 µL cold cell lysis buffer (for 300 µL cell lysis buffer: add 291 µL cell resuspension buffer (500 µL tris-HCl, pH7.5; 125 µL 4M NaCl, 150 µL 1M MgCl2, 49.25 ml autoclaved water), 3 µL 10% NP40 (Roche 11332473001), 3 µL 10% Tween-20 (Roche 11332465001), freshly add 3 µL 1:1 water diluted digitonin (Promega G9441)) was added, and mixed for about 10 times, and samples were placed on ice. After 3 min, 1000 µL wash buffer (990 µL resuspension buffer, 10 µL 10% Tween-20) was added, and samples were centrifugated at 500 RCF for 10 min at 4 °C. The clear supernatants were carefully removed with the pellets intact, then 50 µL tagmentation mixture (25 µL tagmentation buffer (Illumina 15027866), 16.5 µL PBS, 0.5 µL Tween-20, 0.5 µL 1:1 water diluted digitonin, 2.5 µL Tn5 transposase (Illumina 15027865), 5 µL H_2_O) was added to each sample, and placed in a preheated 37 °C thermomixer set to 1000 rpm. After 30 min, the samples were placed on ice, and immediately followed by DNA extraction with MinElute kit (Qiagen). DNA samples were kept at a -80 °C freezer before library preparation. ATAC-seq samples for human primary hepatocytes were performed according to the procedures described above on about 50,000 freshly thawed primary hepatocytes.

### ATAC-seq, and analyses

Library were prepared based on previous method(Buenrostro et al., 2013). After amplifying ATAC-seq library of the specific cycles based on q-PCR measurements, the libraries were double-side selected (0.5-1.3x) with SPRI beads (Beckman), followed by 1x selection to remove primer dimers and sequenced on a Hi-seq 2500 equipment with 60 bp paired-end sequencing.

After removing adaptors with cutadapt (cutadapt -a CTGTCTCTTATA), ATAC-seq results were aligned to Hg38 with bowtie2 (using “--very-sensitive --no-discordant -X 2000”). The generated SAM filers were converted to BAM files, sorted, and indexed using samtools. Bigwig files were generated from BAM files with bamCoverage (-of bigwig), and peak visualizations were performed in a customized session of genome browser(Kent et al., 2002). The total number of reads were obtained with “samtools view -c”, and reads mapped to mitochondrial genomes were removed with “samtools view | grep -v chrM”. PRC duplicates were removed with picard tools (“MarkDuplicates REMOVE_DUPLICATES=true VALIDATION_STRINGENCY=LENIENT”), and properly paired reads were obtained with samtools (“samtools view -h -b -F 1804 -f”). Then reads were shifted (+4 for the forward strand, -5 for the reverse strand) by deeptools (“alignmentSieve --numberOfProcessors 8 –ATACshift”), and blacklist (hg38.blacklist.bed.gz) were used to filter unspecific reads. Post alignment quality controls were performed with ATACseqQC. TSS enrichment scores were calculated with Encode ATAC-seq pipeline (caper run atac.wdl). Fragment length were visualized with Picard CollectInsertSizeMetrics or deeptools bamPEFragmentSize, and reads enrichments were visualized with ngs.plots. ATAC-seq peaks were called with macs2(Zhang et al., 2008) (macs2 callpeaks -f BAMPE -q 0.001 --nomodel -g hs --keep-dup all --cutoff-analysis), and fraction of reads in peaks (FRiP) were performed with BEDTools(Quinlan and Hall, 2010) and SAMtools(Li et al., 2009) (bedtools sort -i peaks.narrowPeak| bedtools merge -i stdin | bedtools intersect -u -nonamecheck -a /final.shifted.bam -b stdin -ubam | samtools view -c). Motif analyses were performed using homer(Heinz et al., 2010) (findMotifsGenome.pl hg38 -size 200 -mask). The analyses of overlapping between THRB-FLAG ChIP-seq (peaks were called from GSM2534017 using the ChIP-seq analysis methods described below) and ATAC-seq peaks were performed using BEDTools.

### Chromatin accessibility measurements with ATAC-seq library and q-PCR

Control PSC-hepatocytes or T3 treated PSC-hepatocytes were tagmented, and tagemented DNA samples were purified and subjected to library preparation as described above. The amplified DNA were purified with MinElute purification kit (Qiagen), and DNA samples (about 0.5-1ng DNA for each sample) were subjected to q-PCR analysis with the following primers identified with ATAC-seq peaks.

A chromatin accessible peak at the *ACTB* locus: F: GCAAAGGCGAGGCTCTGTG; R: CCGTTCCGAAAGTTGCCTTTTATG.

A chromatin accessible peak at the 3’ of *AFP* locus: F: TTGAGGGAACGAAAGGGTGG; R: TTGCCCATGCTCCGTATCTC.

The proximal enhancer of the *CYP3A4* locus: F: CTTGCTGACCCTCTGCTTTCC; R: CTCATGAGGCTGCTTGAACCG.

A chromatin accessible peak encompassing the rs12740374 SNP: F: GTTTGCTCAGTTGCTGACCCAAAAG; R: CCACATCACAGCAAAGAAGCGCAAC. The delta cT valued (cT-*ACTB* cT) were used to compare control samples and T3 treated samples to quantify relative chromatin accessibility.

### Formaldehyde crosslinking, chromatin sonication, H3K27Ac ChIP-seq, and analysis

Crosslinking and ChIP-seq were performed as described (Lee et al., 2006) with modifications described below. Formaldehyde buffer (11% formaldehyde (Tousimis 1008A), 0.5 mM pH8 EGTA, 1 mM pH8 EDTA, 100 mM NaCl, 50 mM pH7.5 HEPES-KOH) was added to cell culture plate at 1:10 ratio to PSC-hepatocytes (2D) from about 4 6-well plates. Cells were crosslinked with for 10 min at room temperature followed by 5 minutes quenching with glycine at room temperature. Cells were collected by scraping and centrifuged for 5 min at 800 RCF at 4°C. After PBS wash twice, the cell pellets were flash frozen in liquid nitrogen and stored in a -80 °C freezer until sonication.

PSC-hepatocytes spheroids were resuspended in 10 ml medium in a 15 ml cornicle tube, and 1 ml crosslinking solution was added, and the tube was put on a 70 rpm horizontal rocker for 10 min at room temperature. Then 0.5 ml 2.5 M glycine solution was added, and quenched for 5 min by putting on the shaker with the same setting. Then organoids were pelleted by 800 RCF centrifugation for 2 min at 4°C, and solutions were removed. The organoids pellets were moved to a 1.5 ml tube, washed with 4 °C PBS, and pelleted by 800 RCF centrifugation for 5 min at 4°C, and repeated wash once. Then PBS was removed and the crosslinked cell pellets were snap frozen in liquid nitrogen, and stored in -80 °C freezer. Primary hepatocytes were crosslink similarly.

Crosslinked material was resuspended in 0.1% SDS (50mM Tris-HCl (pH 8), 10mM EDTA) and sonicated for 5min with a Covaris E220 instrument (5% duty cycle, 140 Peak Incident Power, 200 cycles per burst, 1ml AFA Fiber milliTUBEs). Soluble chromatin (5 µg) is immunoprecipitated with Protein A/G Dynabeads (Thermo Fisher Scientific, 10002D, 10004D) and 10μg of H3K27ac (Diagenode, C15410196). ChIP-seq libraries were constructed using Accel-NGS 2S DNA library kit from Swift Biosciences. Fragments of the desired size were enriched using AMPure XP beads (Beckman Coulter). Libraries were sequenced on an Illumina NovaSeq 6000 instrument with 50 bp paired-end sequencing.

ChIP-seq datasets were aligned using bowtie (Langmead et al., 2009) to the hg38 reference genome using only the standard chromosomes (i.e. chr1 through chr19, chrX, chrY and chrM) with parameter –k 1–m1 –best and –l set to read length.

Peaks were called using MACS (Zhang et al., 2008) with parameters -p 1e-9 –keep-dup=auto and input control. These files were subsequently normalized to the millions of mapped reads and displayed in the UCSC Genome Browser (Kent et al., 2002). Peaks from H3K27Ac ChIP-Seq were used for super-enhancer analyses were performed suing ROSE (Whyte et al., 2013) (https://bitbucket.org/young_computation/rose) with parameters –s 12500 –t 2000. Stitched enhancers were assigned to the single expressed RefSeq gene whose transcription start site was nearest the center of the stitched enhancer.

### Enrichment analysis of liver-disease associated SNPs

Stratified LD score regression(Furuyama et al., 2019) was used to summary statistics from 10 GWAS studies – ALP, ALT, AST, and GGT(Chambers et al., 2011); HDL, LDL, TC and TG(Willer et al., 2013); T2D and T2D stratified by BMI(Mahajan et al., 2018) combined with chromatin data from the tissue samples of the Roadmap Epigenomics and ENCODE projects (Consortium, 2012; Roadmap Epigenomics et al., 2015). Analyses were performed as described (Furuyama et al., 2019), using LDSC v1.0.1 to compare cell types in a model also containing the full baseline model v1.2 annotations.

### Quantification of intracellular urea

Urea quantification was performed with a urea measurement kit (abcam-ab83362). CellTiter-Glo 3D Cell Viability Assay (Promega G9681) were used for normalization.

### Mice and splenic injection of human PSC-differentiated hepatocyte-like cells

Immunodeficient NOD.Cg-*Prkdc^scid^ Il2rg^tm1Wjl^*/SzJ (NSG) mice were obtained from the Jackson Laboratory and housed in autoclaved cages with autoclaved food and water in environmentally controlled rooms at the Whitehead Institute animal facility. Human PSC differentiated hepatocytes spheroids were washed with PBS, then digested with Accutase in a 37 °C water bath for about 20 min with agitation every 3-5 min. The samples were gently pipetted with a 1 mL pipettor for better dissociation and washed with RPMI-1640 medium with 1% BSA-fragment V. The dissociated cells were counted with Countess, resuspended in RPMI-1640 medium and kept on ice until transplantation. NSG recipient mice (2-3 months) were anesthetized with avertin before lateral incisions were made to expose the spleens. About 0.5-1 million cells in about 100 µL RPMI-1640 medium were injected to a spleen of with a 1 ml syringe and a 23-gauge needle. Then the peritoneum and skin were closed with suture and wound clips respectively. The transplanted mice were examined 3 days after transplantation. At 1 month or 6 months post transplantation, transplanted mice were sacrificed and liver tissues were fixed in 10% buffer neutralized formalin solution overnight at 4 °C before tissue processing and paraffin embedding. All mice experiments were in accordance with the protocols approved by the Animal Research Regulation Committee at the Whitehead Institute and guidelines from the Department of Comparative Medicine at Massachusetts Institute of Technology.

### Immunofluorescence staining

Five-micron paraffin slices from FFPE mouse tissue samples were generated with a microtome (Leica) and attached to glass slides. The antigen retrieval and immunofluorescence staining procedures were performed similarly as described above.

Human PSC differentiated hepatocytes spheroids were washed with PBS, and fixed in freshly prepared 4% PFA for 2 hours at room temperature on a rocking platform, followed by wash with PBS twice. Samples were embedded in paraffin followed by tissue processing describe above. The thin sections were deparaffinized followed by antigen retrieval, and blocked with PBS with 0.1% Tween, and 5% donkey serum for 1 hour at room temperature, and incubated with primary antibodies overnight at 4 °C (1:100 mouse anti CYP3A4 antibody, Life Technologies MA5-17064, 1:100 goat anti Albumin antibody, Bethyl A80-229A).

For wholemount staining, the spheroids were fixed as above, followed by incubation in PBS for 20 min, then PBS with 0.5% Tx100 for 20 min. Afterwards, the samples were blocked in PBS with 0.1% Tween and 5% donkey serum for 1 hour, followed by overnight incubation with primary antibody overnight at 4 °C (1:100 rabbit-anti-HNF4A Cell signaling 3113S; 1:250 mouse anti-AFP antibody, Sigma A8452; 1:100 mouse anti CYP3A4 antibody, Life Technologies MA5-17064, 1:100 goat anti Albumin antibody, Bethyl A80-229A). The washing, secondary antibody incubation, DAPI staining, mounting and imaging steps were similar as described above.

### Plasmids

The *CYP3A4*proximal enhancer was predicted based on ATAC-seq datasets. E-crispr (http://www.e-crisp.org/E-CRISP/) was used to design sgRNAs targeting the boundaries of *CYP3A4* enhancer. To generate sgRNA constructs targeting *CYP3A4* proximal enhancer, the annealed products of CACCGGCACATGGTAAACACTAAGA and AAACTCTTAGTGTTTACCATGTGCC were ligated to BbsI linearized Cas9-cherry plasmid to generate 330-cherry-*CYP3A4*-enhancer-L-sgRNA. Similarly, the annealed products of CACCGGAAACTCATGTCCCAATTAA and AAACTTAATTGGGACATGAGTTTCC were ligated to BbsI linearized Cas9-BFP to generate 330-BFP-*CYP3A4*-enhancer-R-sgRNA.

The larger fragment from restriction digestion of pAAVS1-PDi-CRISPRn (Addgene 73500) using AgeI and PacI was ligated to annealed DNA from two oligoes: TAACTTAGGTACCATCCTGCAGGATGGCCGGCCATA and CCGGTATGGCCGGCCATCCTGCAGGATGGTACCTAAGTTAAT. Then the ligated plasmid was digested with KpnI and ClaI, and the dCas9-VP64 (from Addgene 61422) and MS2-p65-HSF1 (from Addgene 61423) fragments were inserted to generate a doxycycline (DOX) inducible CRISPR activation *AAVS1* targeting construct.

To generate sgRNA construct targeting *PDX1* promoter for gene activation, the annealed products of CACCGGCAGGTGCTCGCGGGTACCT and AAACAGGTACCCGCGAGCACCTGCC were ligated to BsmBI linearized lenti sgRNA(MS2)_zeo backbone (Addgene 61427). Similarly, the annealed products from CACCGGATTCACCTGGGGTCAACAC and AAAGTGTTGACCCCAGGTGAATCC were ligated to BsmBI linearized and purified lenti sgRNA_zeo backbone (Addgene 61427) to target proximal enhancer of *CYP3A4*.

### Statistics

Data are expressed as the mean ± s.e.m. or mean ± standard errors, as indicated in the figure legends with *n* and statistical tests. All data are representative of at least two independent experiments (except for ATAC-sequencing). Statistical analyses were performed using Prism 8 software (GraphPad), and statistical significance was reached when *P*≤0.05.

### Data availability

DNA constructs reported in this study are available from Addgene (plasmid ID: 176836-176841). Genetically modified cells used in this study will be available from the corresponding author with an MTA. Next generation sequencing (NGS) data of this study were submitted to the Gene Expression Omnibus (GSE185360).

## Acknowledgments

We thank Drs. S Henikoff and F. Zhang for sharing research reagents; Dr. S. Markoulaki, and R. Flannery for assistance; Drs. P. Sharp, U. Stilz, K. Little, B. Finan, C. Ivashchenko, R, Jing, members of the R.J. laboratory, R.A.Y laboratory, and L.G.G laboratory for discussions; Dr. B. Yuan (Bioinformatics and Research Computing Group of the Whitehead Institute) for assisting with the bioinformatics analysis; W. Salmon (W.M. Keck Biological Imaging Facility of the Whitehead Institute) for assistance with fluorescence imaging; P. Wisniewski and P. Autissier (Whitehead Institute) for assisting with cell sorting; members of the Genome Technology Core at the Whitehead Institute for assisting with sequencing experiments; E Spooner and E. Dudek (Whitehead Institute Proteomics Facility) for helping with mass-spectrometry experiments. This study was supported by a collaborative research agreement from Novo Nordisk, and NIH grant 1R01-NS088538 (to R.J.).

## Author contributions

R.J., R.A.Y., L.G.G., and H.W.L. supervised the research. R.J. and H.M. wrote the manuscript. H.M., E.Z., Y.G., P.C., S.S., and D.F. performed experiments. H.M., P.T., Y.G., M.B., A.D., and B.A. analyzed results. J.F.J. provided advice and edited the manuscript. C.G.E edited the manuscript. T.L. provided advice. L.G. provided primary hepatocytes.

