## Supplemental_figures for "A nuclear receptor facilitates differentiation of human PSCs into more mature hepatocytes"

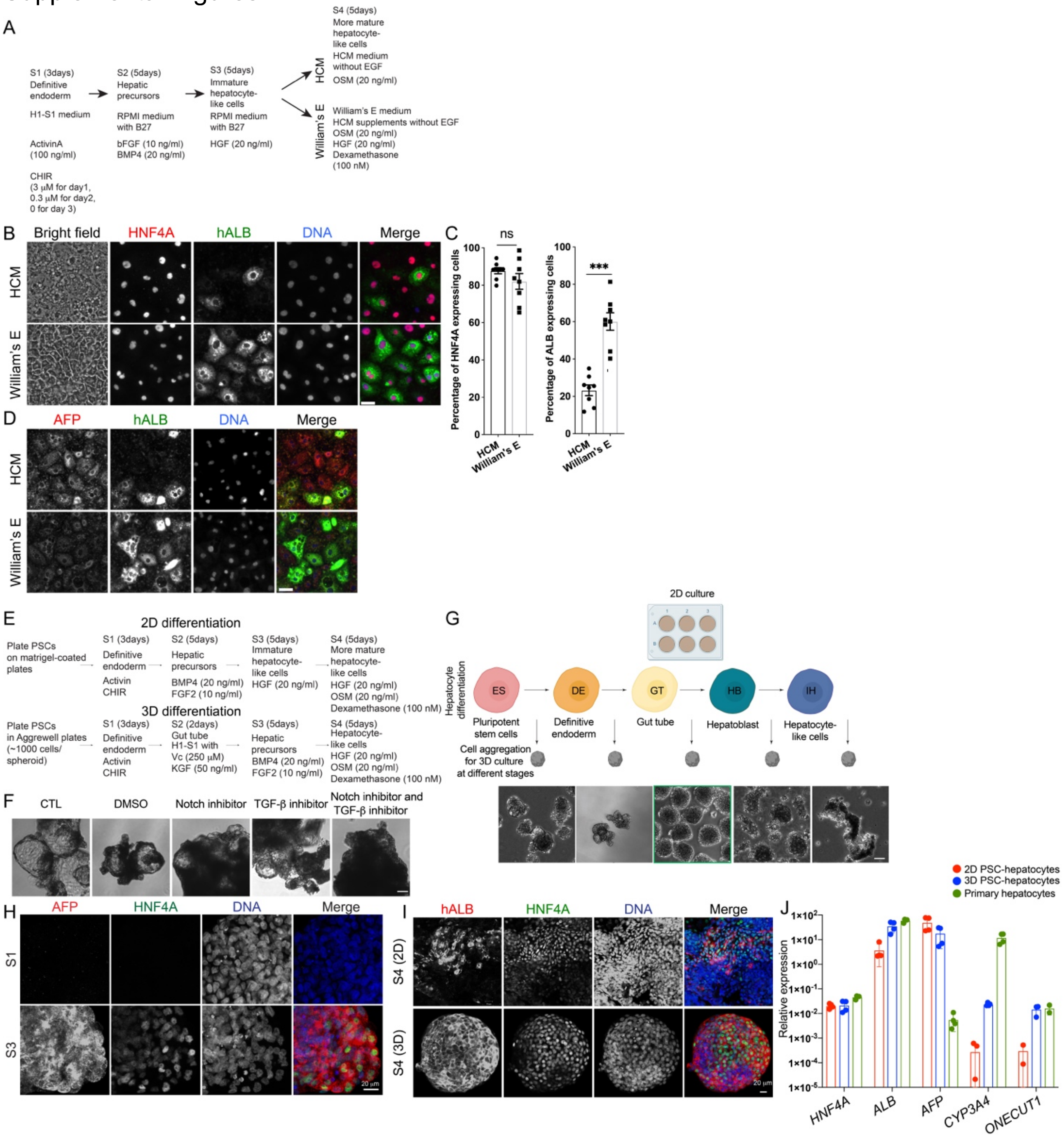


**Supplemental Figure. 1. Development of a 3D culture system for hepatocytes differentiation from human PSC.**

(A) Hepatocytes differentiation protocols in 2D using HCM medium (top panel) or William's E medium (bottom panel).

(B) Representative brightfield and immunofluorescence micrographs of differentiated hepatocyte-like cells with HCM medium (top panel) and William's E medium (bottom panel).

(C) Quantification of the ratio of HNF4A expressing cells (left panel) and human albumin (hALB) expressing cells (right panel) from (B). Plotted data are mean ± s.e.m., *n*=8 regions from 3 experiments. *** denotes *P*<0.001, two-sided Student's *t*-test.

(D) Representative immunofluorescence of expression of fetal hepatocyte marker AFP and adult hepatocyte marker hALB. (a) Comparison between 2D differentiation (top panel) and an initial 3D differentiation protocol (bottom panel).

(E) Comparison between 2D differentiation (top panel) and an initial 3D differentiation protocol (bottom panel).

(F) Addition of Notch inhibitor and TGFb inhibitor suppressed spheroids with cystic morphology.

(G) Optimization of the stages starting cell re-aggregation and spheroid culture.

(H) Representative immunofluorescence micrograph showed expression of AFP and HNF4A after S3.

(I) Representative immunofluorescence micrographs with anti-ALB and anti-HNF4A antibodies on 2D PSC-hepatocytes (top panel) and 3D PSC-hepatocytes (bottom panel).

(J) Gene expression of *HNF4A*, *ALB*, *AFP*, *CYP3A4*, and *ONECUT1* in PSC-hepatocytes differentiation in 2D (red) and 3D (blue), and uncultured primary hepatocytes (green). Plotted data are mean ± standard error.

Scale bars in (B) and (D): 10 μm, scale bars in (F) and (G): 50 μm. Scale bars in (H) and (I): 20 μm.

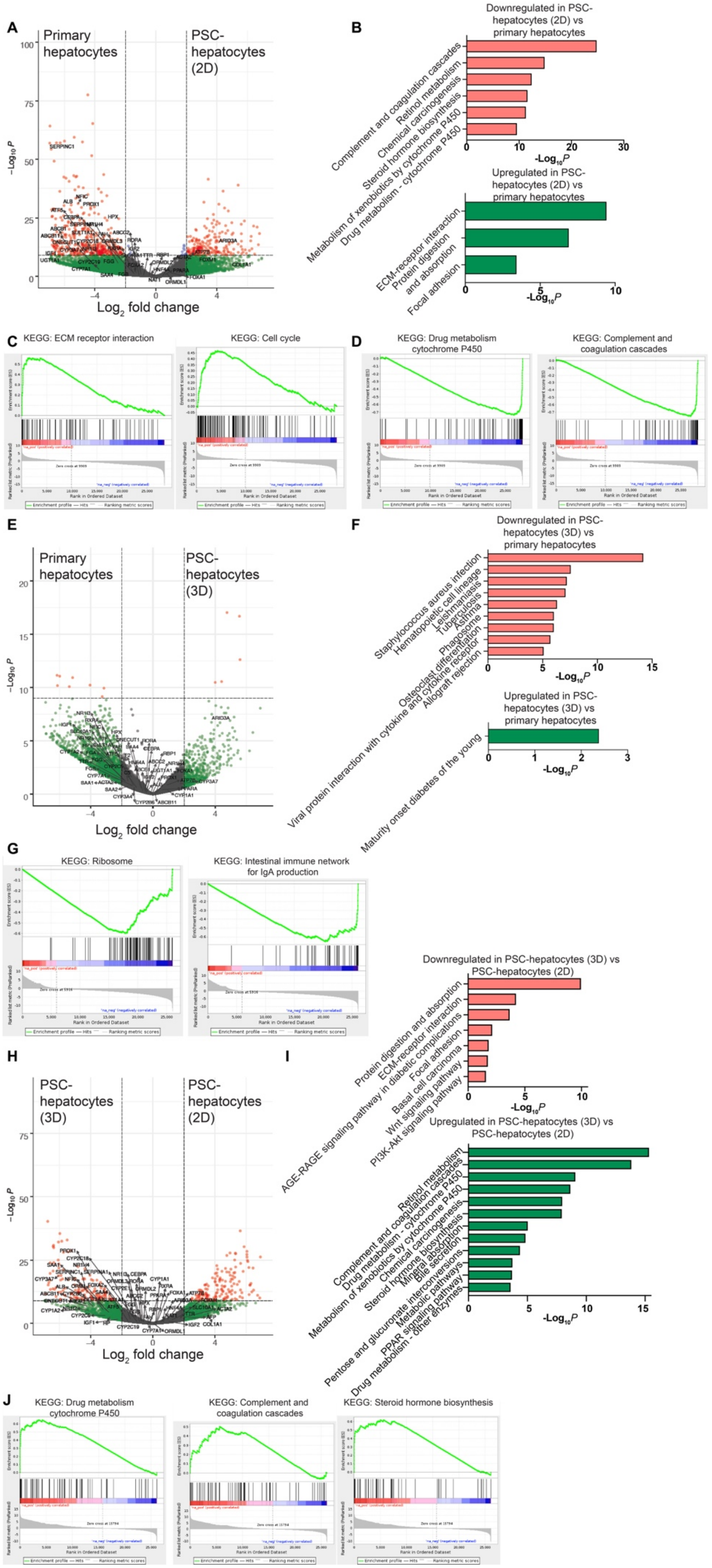
A
Primary hepatocytes
PSC-hepatocytes (2D)
$-Log_{10}P$
Log2 fold change
B
Downregulated in PSC-hepatocytes (2D) vs primary hepatocytes
Complement and coagulation cascades
Retinol metabolism
Chemical carcinogenesis
Steroid hormone biosynthesis
Metabolism of xenobiotics by cytochrome P450
Drug metabolism - cytochrome P450
Upregulated in PSC-hepatocytes (2D) vs primary hepatocytes
ECM-receptor interaction
Protein digestion and absorption
Focal adhesion
C
KEGG: ECM receptor interaction
KEGG: Cell cycle
D
KEGG: Drug metabolism cytochrome P450
KEGG: Complement and coagulation cascades
E
Primary hepatocytes
PSC-hepatocytes (3D)
F
Downregulated in PSC-hepatocytes (3D) vs primary hepatocytes
Staphylococcus aureus infection
Hematopoietic cell lineage
Leishmaniasis
Tuberculosis
Asthma
Phagosome
Osteoclast differentiation
Viral protein interaction with cytokine and cytokine receptor
Allograft rejection
Upregulated in PSC-hepatocytes (3D) vs primary hepatocytes
Maturity onset diabetes of the young
G
KEGG: Ribosome
KEGG: Intestinal immune network for IgA production
H
PSC-hepatocytes (3D)
PSC-hepatocytes (2D)
I
Downregulated in PSC-hepatocytes (3D) vs PSC-hepatocytes (2D)
Protein digestion and absorption
ECM-receptor interaction
AGE-RAGE signaling pathway in diabetic complications
Focal adhesion
Basal cell carcinoma
Wnt signaling pathway
PI3K-Akt signaling pathway
Upregulated in PSC-hepatocytes (3D) vs PSC-hepatocytes (2D)
Retinol metabolism
Complement and coagulation cascades
Drug metabolism - cytochrome P450
Metabolism of xenobiotics by cytochrome P450
Chemical carcinogenesis
Steroid hormone biosynthesis
Mineral absorption
Bile secretion
Pentose and glucuronate interconversions
Metabolic pathways
PPAR signaling pathway
Drug metabolism - other enzymes
J
KEGG: Drug metabolism cytochrome P450
KEGG: Complement and coagulation cascades
KEGG: Steroid hormone biosynthesis

**Supplemental Figure. 2. Additional comparison of RNA-seq results from primary hepatocytes, 2D, and 3D PSC-hepatocytes.**

(A) Volcano plots of RNA-seq between primary hepatocytes and 2D PSC-hepatocytes. Green spots denotes genes with more than 4 folds changes. Purple spots denote $P<10^{-9}$. Red spots denote genes with both larger than 4 folds changes and $P<10^{-9}$. Grey dots denote genes that are not significantly different.

(B) KEGG pathway analysis of pathways enriched in genes downregulated in 2D PSC-hepatocytes (top panel in red) and genes upregulated in 2D PSC-hepatocytes (bottom panel in green).

(C) GESA of pathways significantly enriched in 2D PSC-hepatocytes compared to primary hepatocytes.

(D) GESA of pathways significantly enriched in primary hepatocytes compared to 2D PSC-hepatocytes.

(E) Volcano plots of RNA-seq between primary hepatocytes and 3D PSC-hepatocytes.

(F) KEGG pathway analysis of pathways enriched in genes downregulated in 3D PSC-hepatocytes (top panel in red) and genes upregulated in 3D PSC-hepatocytes (bottom panel in green).

(G) GESA of pathways significantly enriched in primary hepatocytes compared to 3D PSC-hepatocytes.

(H) Volcano plots of RNA-seq between 3D PSC-hepatocytes and 2D PSC-hepatocytes.

(I) KEGG pathway analysis of pathways enriched in genes downregulated in 3D PSC-hepatocytes compared to 2D PSC-hepatocytes (top panel) and genes upregulated in 3D PSC-hepatocytes compared to 2D PSC-hepatocytes (bottom panel).

(J) GESA of pathways significantly enriched in 3D PSC-hepatocytes compared to 2D PSC-hepatocytes.

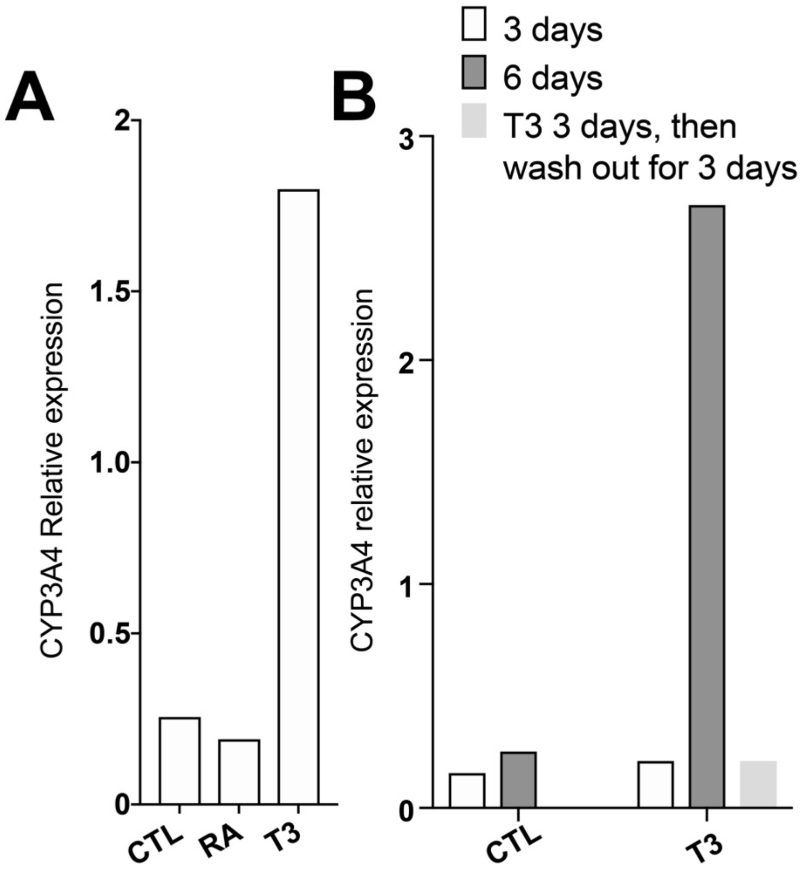


**Supplemental Figure 3. Characterization of the effects of retinoic acid (RA) and T3 on *CYP3A4* expression in PSC-hepatocytes.**

(A) Q-RT-PCR analysis of control PSC-hepatocytes, or PSC-hepatocytes treated with 20 nM RA or 3 μM T3 for 1 week.

(B) Q-RT-PCR analysis of requirement of maintenance of *CYP3A4* expression by T3.

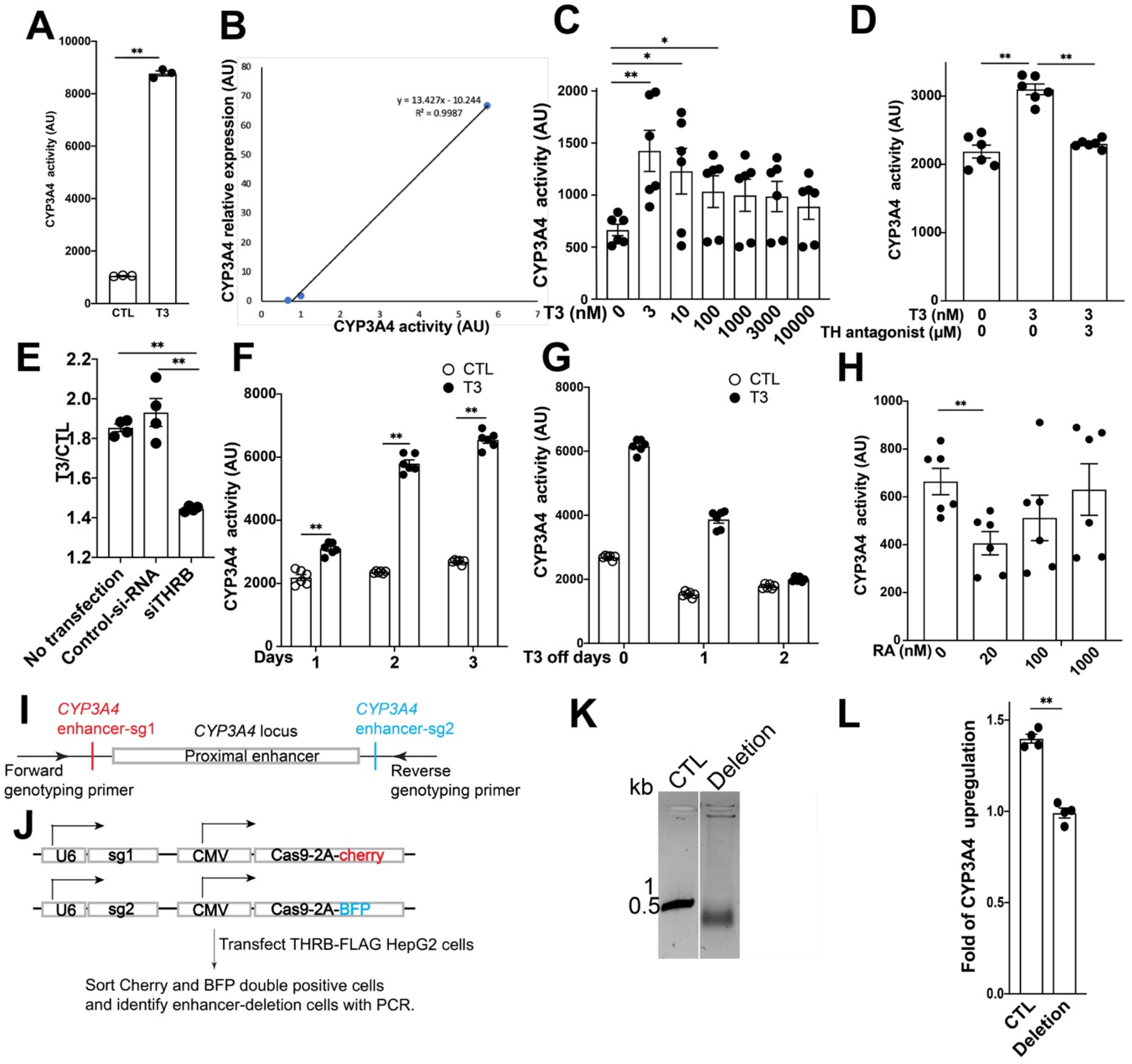


**Supplemental Figure. 4. T3 regulated CYP3A4 in THRB-FLAG-HepG2 cells.**

(A) THRB-FLAG HepG2 cells were cultured in William's E medium (CTL) or William's E medium with T3 for approximately 2 weeks before CYP3A4 activity were measured. Data were plotted as mean ± s.e.m., *n*=3.

(B) Correlation between *CYP3A4* expression and CYP3A4 activity in HepG2-THRB-FLAG cells.

(C) The effects of different concentration of T3 on CYP3A4 activity of THRB-FLAG-HepG2 cells. Data were plotted as mean ± s.e.m., *n*=6. ** denotes *P*<0.01, * denotes *P*< 0.05, two-sided Student's *t* test.

(D) Upregulation of CYP3A4 activity by T3 was reduced by thyroid hormone antagonist. Data were plotted as mean ± s.e.m., $n$=6. ** denotes $P$<0.01, two-sided Student's $t$ tests.

(E) THRB-FLAG-HepG2 cells were transfected with control-si-RNA or esi-RNA targeting *THRB*, followed by incubation with T3-containing medium, and CYP3A4 activity measurement. The ratio of CYP3A4 activity in T3 containing medium over control medium were plotted as mean ± s.e.m., $n$=4. ** denotes $P$<0.01, two-sided Student's $t$ test.

(F) THRB-FLAG-HepG2 cells were cultured in T3-containing medium or control medium for different days, followed by CYP3A4 activity measurements. Data were plotted as mean ± s.e.m., $n$=6. ** denotes $P$<0.01, * denotes $P$< 0.05, two-sided Student's $t$ test.

(G) THRB-FLAG-HepG2 cells cultured in T3-containing medium were changed to control medium for different days, followed by CYP3A4 activity measurements. Data were plotted as mean ± s.e.m., $n$=6.

(H) Effects of retinoic acid (RA) on CYP3A4 activity of THRB-FLAG-HepG2 cells. Data were plotted as mean ± s.e.m., $n$=6. ** denotes $P$<0.01, * denotes $P$< 0.05, two-sided Student's $t$ test.

(I) Experimental design to target *CYP3A4* proximal enhancer boundaries with *CYP3A4* enhancer sg1 and *CYP3A4* enhancer sg2.

(J) THRB-FLAG HepG2 cells were transfected with 2 plasmids (Cas9-2A-cherry-sg1 and Cas9-2A-BFP-sg2), and cherry/BFP double positive cells were isolated to identify *CYP3A4* enhancer deletion clones with PCR.

(K) Identification of *CYP3A4* proximal enhancer deletion cells with genotyping primer illustrated in (H).

(L) CYP3A4 activity assays showed reduced CYP3A4 activity upregulation by T3 in *CYP3A4* proximal enhancer deleted THRB-FLAG HepG2 cells compared to control cells. Data were plotted as mean ± s.e.m., $n$=4. ** denotes $P$<0.01, two-sided Student's $t$ test.

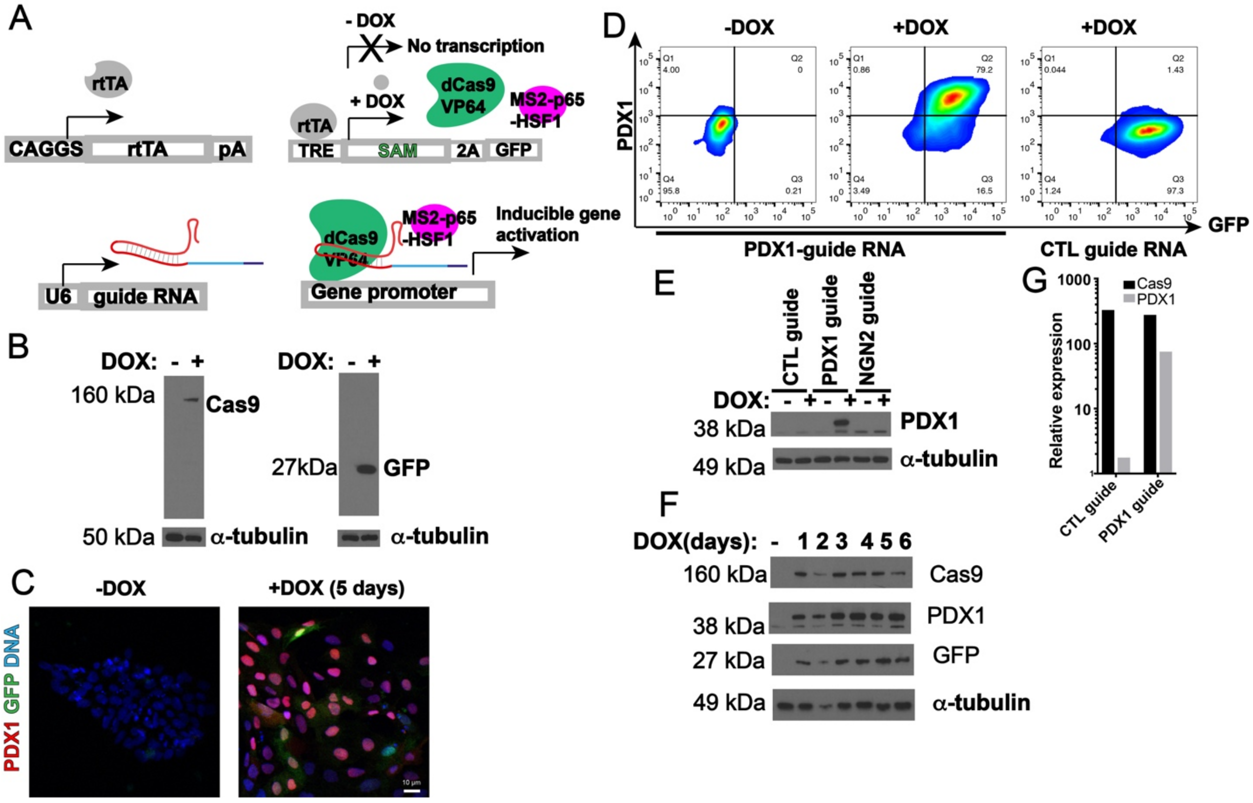


**Supplemental Figure. 5. Characterization of an inducible CRISPR activation system in human PSCs.**

(A) Experimental design to target rtTA expressing cassette and tetracycline response element (TRE)-CRISPR- activation cassette to each allele of the *AAVS1* locus.

(B) Western blotting verification of DOX inducible expression of dCas9 and GFP with anti-Cas9 antibody (left panel) and anti-GFP antibody (right panel).

(C) Immunofluorescence staining showing DOX inducible expression of PDX1 in inducible CRISPR activation PSCs expressing PDX1-sgRNA.

(D) Flow cytometry analysis of PDX1 expression in inducible CRISPR activation cells expressing *PDX1*-sgRNA (left 2 panels) and control cells (right panel).

(E) Western blotting analysis of PDX1 expression in control inducible CRISPR activation cells, the cells expressing PDX1-sgRNA, and the cells expressing NGN2 sgRNA.

(F) Western blotting analysis of dynamics of PDX1 induction in inducible CRISPR activation cells expressing *PDX1* sgRNA.

(G) Q-RT-PCR analysis of *PDX1* expression in CRISPR activation cells.

The scale bar in (C): 10 μm.

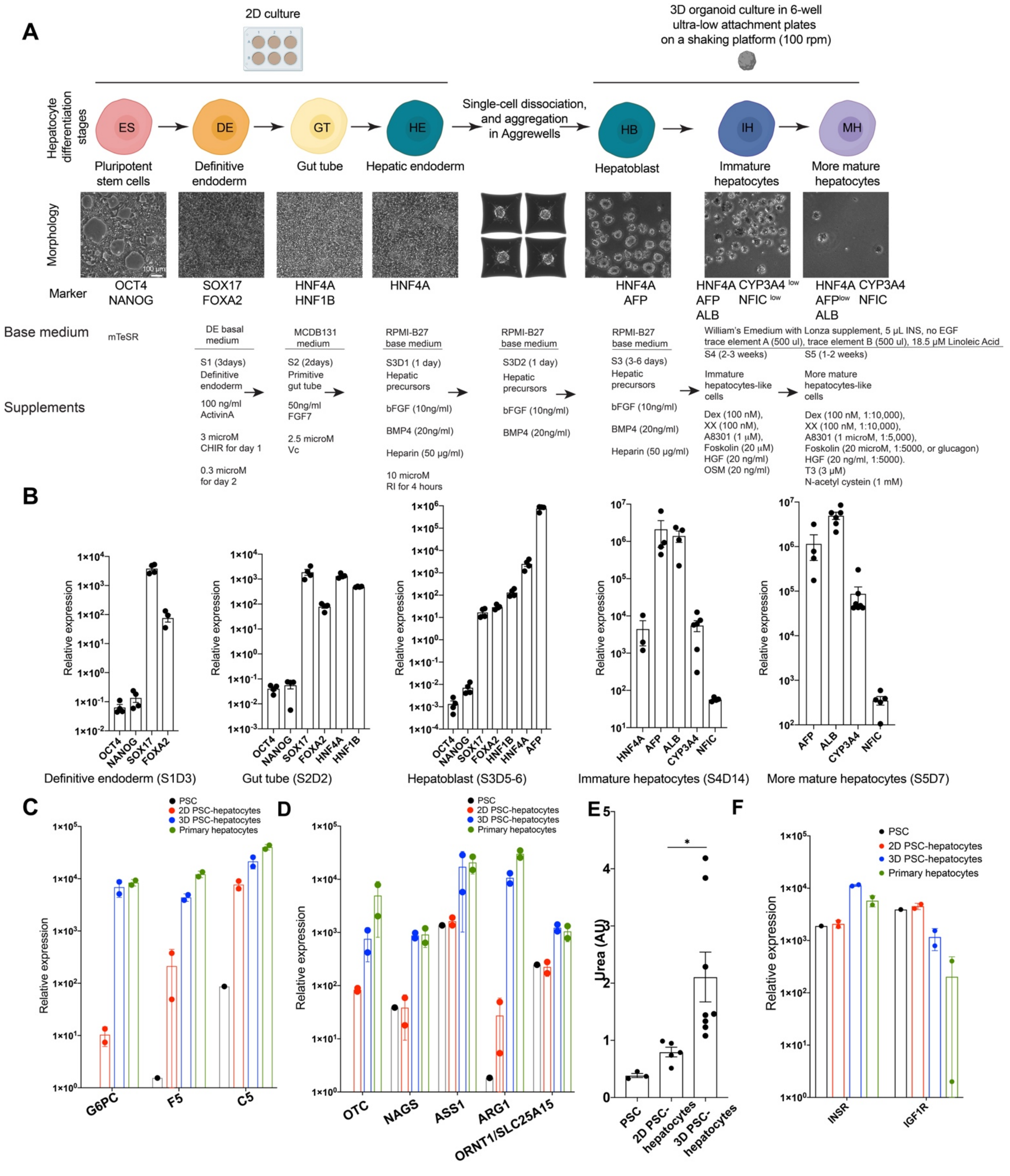


**Supplemental Figure. 6. Stage-specific gene expression during spheroid-based hepatocytes differentiation and functional characterization of PSC-hepatocytes.**

(A) The protocol of spheroid-based hepatocytes differentiation.

(B) Stage specific expression of key genes during hepatocyte differentiation. Plotted data are mean ± s.e.m., *n*=4.

(C) Expression of selected genes mutation of which are associated with genetic disease. Plotted data are mean ± standard error, *n*=2.

(D) Expression of urea cycle genes. Plotted data are mean ± standard error, *n*=2.

(E) Quantification of urea synthesis in PSC, 2D, and 3D PSC-hepatocytes. Plotted data are mean ± s.e.m., *n*=3 for PSCs, *n*=5 for 2D PSC-hepatocytes (2D), and *n*=8 for 3D PSC-hepatocytes (3D). * denotes $P$<0.05, two-sided Student's $t$ test.

(F) Expression of insulin receptor (*INSR*) and Insulin-like growth factor 1 receptor (*IGF1R*). Plotted data are mean ± standard error, *n*=2.

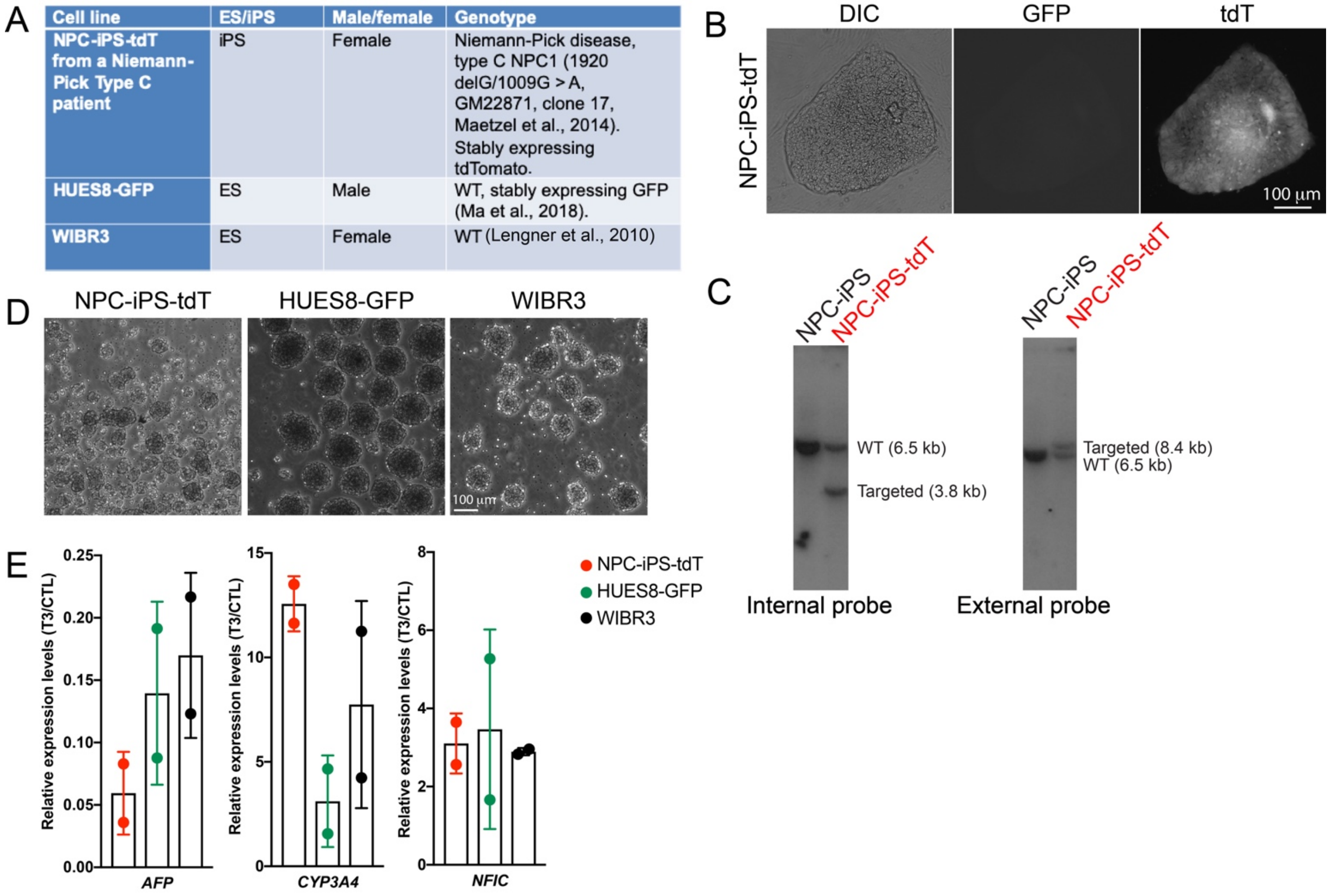

| Cell line | ES/iPS | Male/female | Genotype |
| --- | --- | --- | --- |
| NPC-iPS-tdT from a Niemann-Pick Type C patient | iPS | Female | Niemann-Pick disease, type C NPC1 (1920 delG/1009G > A, GM22871, clone 17, Maetzel et al., 2014). Stably expressing tdTomato. |
| HUES8-GFP | ES | Male | WT, stably expressing GFP (Ma et al., 2018). |
| WIBR3 | ES | Female | WT (Lengner et al., 2010) |



**Supplemental Figure 7. T3 treatment improved gene expression of hepatocytes differentiated from additional human PSCs.**

(A) Information of additional human PSCs tested for hepatocyte differentiation: a tdTomato-expressing iPS from an NPC patient (Maetzel et al., 2014) generated by integration of a tdTomato-expressing cassette in the *AAVS1* locus, a GFP-expressing ES cell line HUES8-GFP (Ma et al., 2018), and a female ES cell line WIRBR3 (Lengner et al., 2010).

(B) Fluorescence microscopy showed expression of tdTomato in NPC-iPS-tdT cells.

(C) Southern blotting characterization of integration of tdTomato expressing cassette to the *AAVS1* locus in NPC-iPS cells.

(D) Morphology of hepatocytes spheroids differentiated from the three human PSC cell lines.

(E) Expression of *AFP*, *CYP3A4*, and *NFIC* when hepatocytes spheroids were changed to T3 containing medium compared to control medium. Plotted data are mean ± standard error, *n*=2.

Scale bars in (B) and (D): 100 μm.
